# Constraint-Based Pan-Cancer Metabolic Modeling Identifies Medium-Robust Biomarkers and Broad-Spectrum Anticancer Targets

**DOI:** 10.64898/2026.11.24.769069

**Authors:** Feng-Sheng Wang, Syuan-Yu Pan, Yuan-Fu Yeh, Kuang-Tse Huang

**Affiliations:** Department of Chemical Engineering, National Chung Cheng University, Chiayi 621301, Taiwan; Research Center for Systems Biology and Tissue Engineering, National Chung Cheng University, Chiayi 621301, Taiwan

**Author notes:** Correspondence (F.S.W.); (K.T.H.).

**Keywords:** Computational systems biology, cancer metabolism, flux balance analysis, hybrid differential evolution, fuzzy multi-objective optimization, anticancer target discovery

## Abstract

**Background:** Cancer metabolic reprogramming varies across tumour types and nutrient environments, complicating the identification of robust metabolic biomarkers and therapeutic targets. We developed a constraint-based pan-cancer framework to identify medium-robust metabolic alterations and broadly effective candidate enzyme targets.

**Methods:** Transcriptomic profiles from 33 TCGA cancer types were integrated with Recon3D to reconstruct cancer- and healthy-tissue-specific genome-scale metabolic models. Parsimonious multi-objective metabolite flow variability analysis was performed under five nutrient conditions to classify differential rates of metabolite synthesis, uptake, and secretion. An extended Anticancer Target Discovery framework incorporating fuzzy multi-objective optimization was employed to evaluate candidate enzyme perturbations based on cancer-cell inhibition, ATP suppression, preservation of healthy-tissue metabolism, and robustness across nutrient conditions.

**Findings:** The models predicted recurrent increases in cardiolipin-related lipid synthesis and aromatic amino acid metabolism, enhanced uptake of essential amino acids and choline, and elevated lactate secretion across multiple cancer–medium combinations. CRLS1 emerged as the most broadly effective predicted enzyme target, satisfying the treatment criteria across all 165 cancer–medium combinations, whereas PGS1 and PTPMT1 exhibited broad but incomplete coverage. These findings highlight mitochondrial phospholipid metabolism as a potential shared vulnerability across cancer types and provide testable candidates for metabolomic and experimental validation.

## 1. Introduction

Metabolic reprogramming is a hallmark of cancer, enabling malignant cells to sustain proliferation, adapt to metabolic and oxidative stress, remodel the tumour microenvironment, and resist therapy. Since Warburg’s description of aerobic glycolysis, our understanding of cancer metabolism has expanded to encompass mitochondrial function, amino acid utilization, nucleotide biosynthesis, lipid metabolism, one-carbon metabolism, and redox regulation [1–7]. Pan-cancer studies have further revealed both shared metabolic alterations across tumour types and context-dependent adaptations influenced by tissue lineage, genetic background, microenvironmental interactions, and nutrient availability [8–12].

Genome-scale metabolic models (GSMMs) provide a mechanistic framework for translating molecular profiles into feasible metabolic phenotypes. Comprehensive human metabolic reconstructions, including Recon, Recon2, Recon3D, and Human1, integrate metabolites, biochemical reactions, subcellular compartments, and gene–protein–reaction associations [13–16]. Combined with flux balance analysis (FBA) and flux variability analysis (FVA), these models enable the characterization of feasible metabolic states, pathway flexibility, and potential metabolic vulnerabilities under defined environmental conditions [17–19]. Accordingly, context-specific GSMMs have been widely applied to investigate cancer cell lines and patient-derived tumours, identify altered metabolic pathways and essential reactions, and predict potential therapeutic targets [20–31].

Despite these advances, identifying robust metabolic biomarkers remains challenging because model predictions can be sensitive to nutrient composition, reconstruction procedures, objective functions, and cancer type. Moreover, changes in gene expression do not necessarily correspond to changes in metabolic fluxes, as pathway activities are constrained by network stoichiometry, substrate availability, and cellular metabolic demands. Constraint-based flux analysis can partially address these limitations by integrating transcriptomic profiles with stoichiometric and environmental constraints, thereby providing a functional interpretation of cancer-associated metabolic alterations.

Metabolite synthesis, uptake, and secretion rates provide complementary indicators of metabolic activity and exchange. However, predicted fluxes should not be directly interpreted as metabolite concentrations, which also depend on transport, consumption, turnover, and compartmentalization. Consequently, model-derived metabolic alterations should be regarded as candidate biomarkers requiring experimental validation. Parsimonious metabolite flow variability analysis (pMFVA) was previously developed to characterize differences in metabolite synthesis and exchange fluxes between cancer- and healthy-tissue-specific models. Extending this approach across cancer types and nutrient environments may facilitate the identification of consistent metabolic alterations while distinguishing medium-robust biomarkers from cancer-specific and nutrient-dependent responses.

Beyond biomarker discovery, identifying effective and selective anticancer targets presents an additional challenge. An ideal metabolic target should suppress cancer-cell growth and ATP production while preserving essential metabolic functions in healthy tissues and maintaining its inhibitory effects across nutrient environments. These competing objectives require a systematic approach to balancing therapeutic efficacy, metabolic selectivity, and robustness. Fuzzy multi-objective optimization provides a suitable framework for addressing such trade-offs by integrating multiple treatment objectives through fuzzy satisfaction criteria. The Anticancer Target Discovery (ACTD) framework combines context-specific GSMMs, constraint-based analysis, and fuzzy multi-objective optimization to prioritize candidate metabolic perturbations [27–31]. Specifically, ACTD evaluates enzyme targets based on their predicted effects on cancer-cell biomass production, ATP synthesis, and metabolic deviations from cancer and healthy reference states. Extending this framework across cancer types and nutrient conditions may enable the identification of broadly effective targets while distinguishing shared metabolic vulnerabilities from tumour-specific or environment-dependent responses.

In this study, transcriptomic profiles from 33 TCGA cancer types were integrated with Recon3D to reconstruct cancer- and healthy-tissue-specific GSMMs, which were subsequently evaluated under five nutrient conditions. We extended pMFVA to parsimonious multi-objective metabolite flow variability analysis (pMOMFVA) to identify candidate metabolic biomarkers exhibiting consistent differential synthesis, uptake, or secretion patterns across nutrient environments. Furthermore, we extended ACTD by incorporating fuzzy multi-objective optimization to prioritize enzyme targets based on predicted cancer-cell inhibition, ATP suppression, preservation of healthy-tissue metabolism, metabolic deviation from cancer-associated states, and robustness across nutrient conditions. Together, these approaches establish an integrated computational framework for identifying shared pan-cancer metabolic alterations, robust candidate biomarkers, and broadly effective anticancer targets.

## 2. Materials and methods

### 2.1. Reconstruction of pan-cancer and healthy-tissue metabolic models

Tissue-specific genome-scale metabolic models (GSMMs) were reconstructed using RNA-sequencing profiles obtained from The Cancer Genome Atlas (TCGA) [32]. The dataset comprised 33 cancer types, many of which included both tumour and matched adjacent-normal samples. Tumour and normal expression profiles were processed separately using a reconstruction pipeline involving quantile normalization, statistical analysis, and classification of gene expression levels [29, 31]. The resulting expression classifications were integrated with gene–protein–reaction (GPR) associations in Recon3D to reconstruct cancer-associated (CA) and healthy-tissue (HT) metabolic models.

Nine cancer types—adrenocortical carcinoma (ACC), diffuse large B-cell lymphoma (DLBC), acute myeloid leukaemia (LAML), lower-grade glioma (LGG), mesothelioma (MESO), ovarian serous cystadenocarcinoma (OV), testicular germ-cell tumour (TGCT), uterine carcinosarcoma (UCS), and uveal melanoma (UVM)—lacked matched adjacent-normal samples, whereas skin cutaneous melanoma (SKCM) contained only one normal sample. For these ten cancer types, the generic Recon3D model was used as the HT reference [15]. Consequently, metabolic comparisons involving these cancer types may reflect differences in tissue specificity and model composition in addition to cancer-associated metabolic alterations.

### 2.2. Identification of candidate metabolic biomarkers

The workflow for identifying candidate metabolic biomarkers is illustrated in Figure 1. Although differential gene-expression (DGE) analysis identifies transcriptomic differences between biological conditions, these changes do not necessarily correspond to differences in feasible metabolic activity [33]. To address this limitation, we extended the analysis to the functional metabolic level by comparing predicted metabolite synthesis, uptake, and secretion rates between CA and HT models. This approach integrates genome-scale metabolic reconstructions with flux-based optimization to characterize cancer-associated metabolic alterations [29, 30].

**Figure 1.**
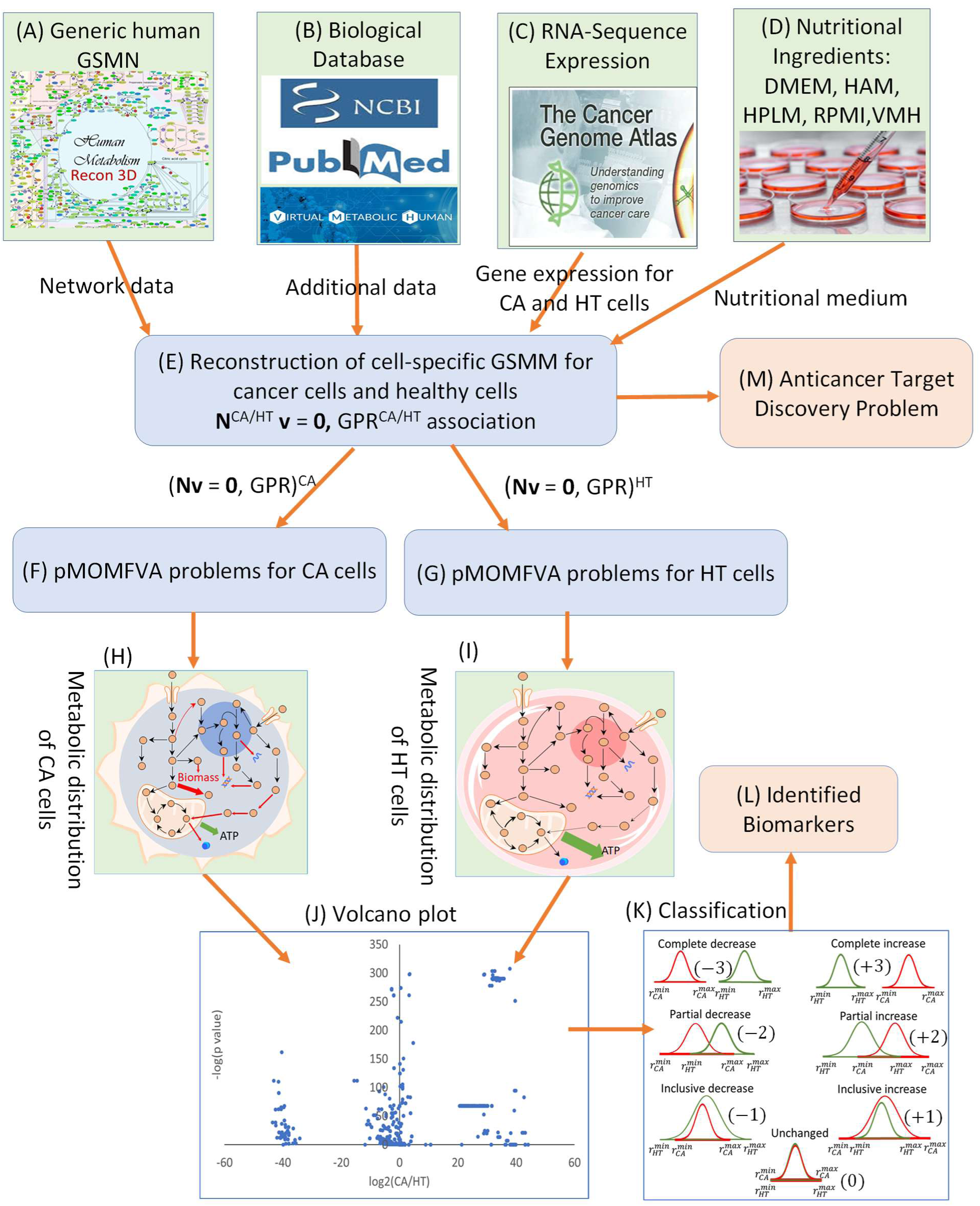
Workflow for candidate metabolic biomarker identification using differential analysis and parsimonious multi-objective metabolite flow variability analysis. (A) Recon3D is used as the generic human metabolic-network template. (B) Biological databases provide context-specific nutrient and pathway information. (C) RNA-sequencing profiles from cancer and corresponding healthy tissues are integrated into the reconstruction procedure. (D) The reconstructed models are evaluated under five nutrient media. (E) Tissue-specific GSMMs and their GPR associations are generated for cancer-associated (CA) and healthy-tissue (HT) conditions. (F,G) pMOMFVA is applied separately to the CA and HT models to determine feasible flux and metabolite-flow ranges. (H, I) Metabolic distributions are obtained for the CA and HT models, respectively. (J) Log-ratios and corresponding *p*-values are calculated for metabolite synthesis and exchange rates, where 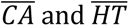 denote the corresponding mean rates. (K) CA and HT ranges are assigned to seven categories according to their direction and degree of overlap. (L) Candidate metabolic biomarkers are selected using the classification scores and statistical criteria. (M) The reconstructed models are subsequently used in the ACTD analysis.

Parsimonious multi-objective metabolite flow variability analysis (pMOMFVA) was applied to determine the feasible ranges of metabolite flow rates under each nutrient condition. The resulting CA and HT ranges were compared according to their relative positions and degree of overlap and classified into seven categories: complete increase (CI), partial increase (PI), inclusive increase (II), unchanged (UC), inclusive decrease (ID), partial decrease (PD), and complete decrease (CD) (Figure 1K). Numerical scores were assigned to each category—CI (+3), PI (+2), II (+1), UC (0), ID (−1), PD (−2), and CD (−3)—and the mean score was calculated across the five nutritional media.

For each metabolite or exchange reaction, the classification scores were averaged across the five nutrient conditions. Candidate medium-robust biomarkers were identified when the absolute average score was at least 2.4 and the corresponding p-value was less than 0.05. Positive average scores indicated higher predicted flow rates in CA models, whereas negative scores indicated lower rates relative to their corresponding HT models. Among the seven categories, CI and CD represented non-overlapping CA and HT ranges and therefore provided the strongest model-based discrimination between cancer and healthy-tissue metabolic states.

### 2.3. Parsimonious multi-objective metabolite flow variability analysis

Five nutrient media—Dulbecco’s modified Eagle medium (DMEM), Ham’s medium (HAM), human plasma-like medium (HPLM), Roswell Park Memorial Institute medium (RPMI), and Virtual Metabolic Human medium (VMH)—were considered to evaluate the robustness of predicted metabolic alterations across different nutrient conditions.

The pMOMFVA framework was formulated as a bilevel optimization problem. At the upper level, the synthesis rate of each metabolite was independently maximized and minimized to determine its feasible range. The lower-level problem comprised two sequential optimization stages: multi-objective flux balance analysis (MOFBA) and uniform flux distribution (UFD). The MOFBA stage determined optimal cellular objective values, whereas the UFD stage identified a parsimonious flux distribution subject to constraints that preserved specified fractions of these optimal values. The overall formulation is presented in Equation (1).

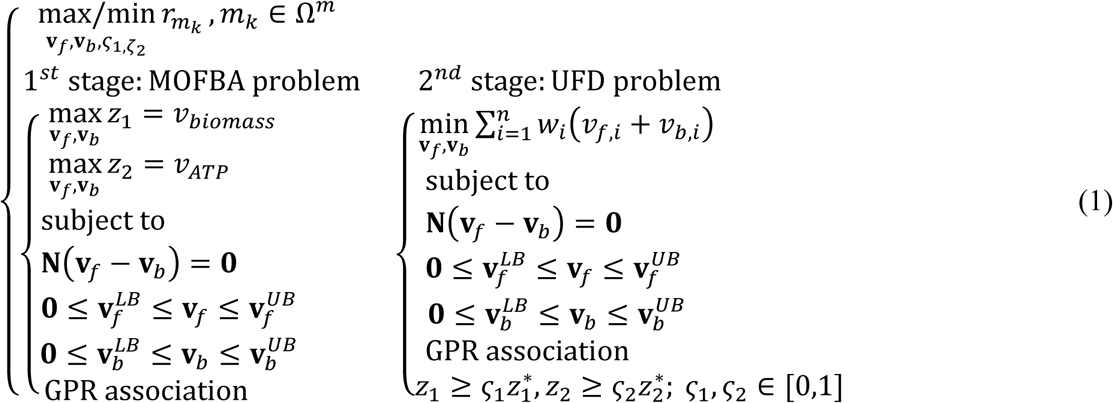

The forward and backward flux vectors, **v**_ƒ_ and **v**_b_, represent the two directional components of each metabolic reaction. The stoichiometric matrix **N** ∈ ℝ^m×n^, contains *m* metabolites and *n* reactions. Steady-state mass balance was imposed through **N**(**v**_ƒ_ − **v**_b_) = **0**, with reaction fluxes constrained by their respective lower bounds 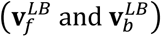 and upper bounds 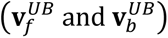. GPR associations were incorporated into the reaction constraints and weighting scheme to account for transcriptomic support.

For CA models, MOFBA simultaneously maximized the biomass growth rate, *v*_biomass_, and ATP synthesis rate, *v*_ATP_, representing cellular proliferative and energetic demands, respectively. In contrast, ATP synthesis alone was maximized for HT models to represent energy maintenance rather than proliferative growth. Because CA and HT models were assigned different cellular objectives, some predicted differences in anabolic fluxes may arise from objective-function specifications in addition to differences in transcriptomic profiles and metabolic network composition.

In the UFD stage, the weighted sum of forward and backward reaction fluxes was minimized to obtain a parsimonious flux distribution:

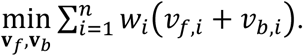

The reaction weights *w*_i_ were assigned according to transcriptomic confidence levels:

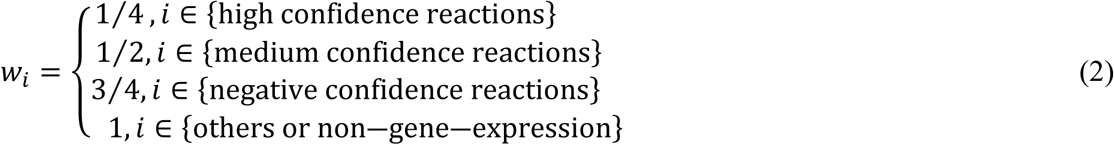

These expression-dependent weights represent relative penalties on reaction fluxes rather than experimentally calibrated kinetic parameters. Reactions with stronger transcriptomic support were assigned lower weights and consequently incurred smaller penalties during flux minimization. This weighting strategy favours parsimonious flux distributions that are more consistent with the underlying gene-expression profiles.

Wang and Zhang [31] previously evaluated the influence of reaction weighting on metabolic flux predictions by comparing hepatocellular carcinoma and healthy-tissue models using parsimonious flux balance analysis with either expression-dependent or uniform reaction weights (*w*_i_ = 1). The predicted directions of metabolic flux changes were assessed against the Warburg hypothesis and established metabolic alterations documented in the literature [5, 6, 61, 62]. Expression-dependent weighting yielded a reported prediction accuracy of 63.9%, compared with 59.1% for uniform weighting, representing an improvement of 4.8 percentage points. These findings suggest that incorporating transcriptomic confidence into flux minimisation can improve the biological consistency of constraint-based metabolic predictions.

For a metabolite *m* in compartment *c,* the compartment-specific synthetic rate was calculated as

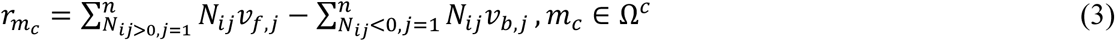

Here, *N*_ij_ denotes the stoichiometric coefficient of metabolite *i* in reaction *j*, whereas *v*_ƒ,j_ and *v*_b,j_ represent the corresponding forward and backward fluxes. The first term accounts for metabolite production through forward reactions, whereas the second represents production through backward reactions. Because the stoichiometric coefficients in the second term are negative, subtraction yields a positive contribution to metabolite production. For metabolites distributed across multiple compartments, the overall synthesis rate was obtained by summing the corresponding compartment-specific rates, as expressed in Equation (4).

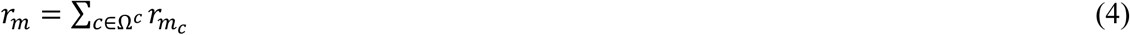

The summation is performed over the set of compartments Ω^c^ containing metabolite *m*.

To accommodate multiple cellular objectives, the first-stage MOFBA problem was reformulated using fuzzy set theory and the priority-based maximizing fuzzy decision (PBMFD) approach. This formulation integrates competing cellular objectives through membership functions while accounting for their relative priorities. The resulting PBMFD–UFD formulation is presented in Equation (5).

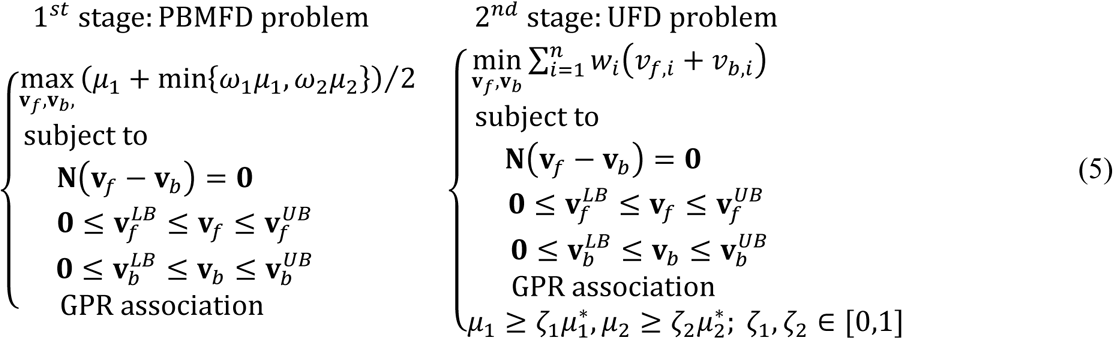

The weighing factor *ω*_i_ represents the relative priority assigned to membership function *μ*_i_. For each maximized objective, the corresponding membership grade was calculated using Equation (6).

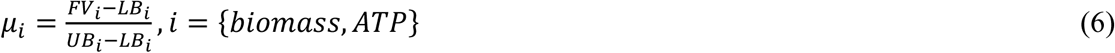

Here, *FV*_i_ denotes the calculated objective *i*, whereas *LB*_i_ and *UB*_i_represent its predefined lower and upper bounds, respectively. These membership grades provide normalized measures of objective satisfaction and are incorporated into the PBMFD formulation to determine the preferred metabolic flux distribution.

### 2.4. Anticancer target discovery

The Anticancer Target Discovery (ACTD) framework, illustrated in Figure 2, was employed to identify candidate enzyme perturbations predicted to inhibit cancer-cell growth and ATP production while preserving essential metabolic functions in the corresponding healthy-tissue models. Candidate targets were evaluated using a fuzzy multi-objective hierarchical optimization framework that integrates cancer-cell inhibition, healthy-tissue viability preservation, and treatment-induced metabolic deviation. These objectives were considered jointly to balance predicted anticancer efficacy against potential adverse metabolic effects on healthy tissues.

**Figure 2.**
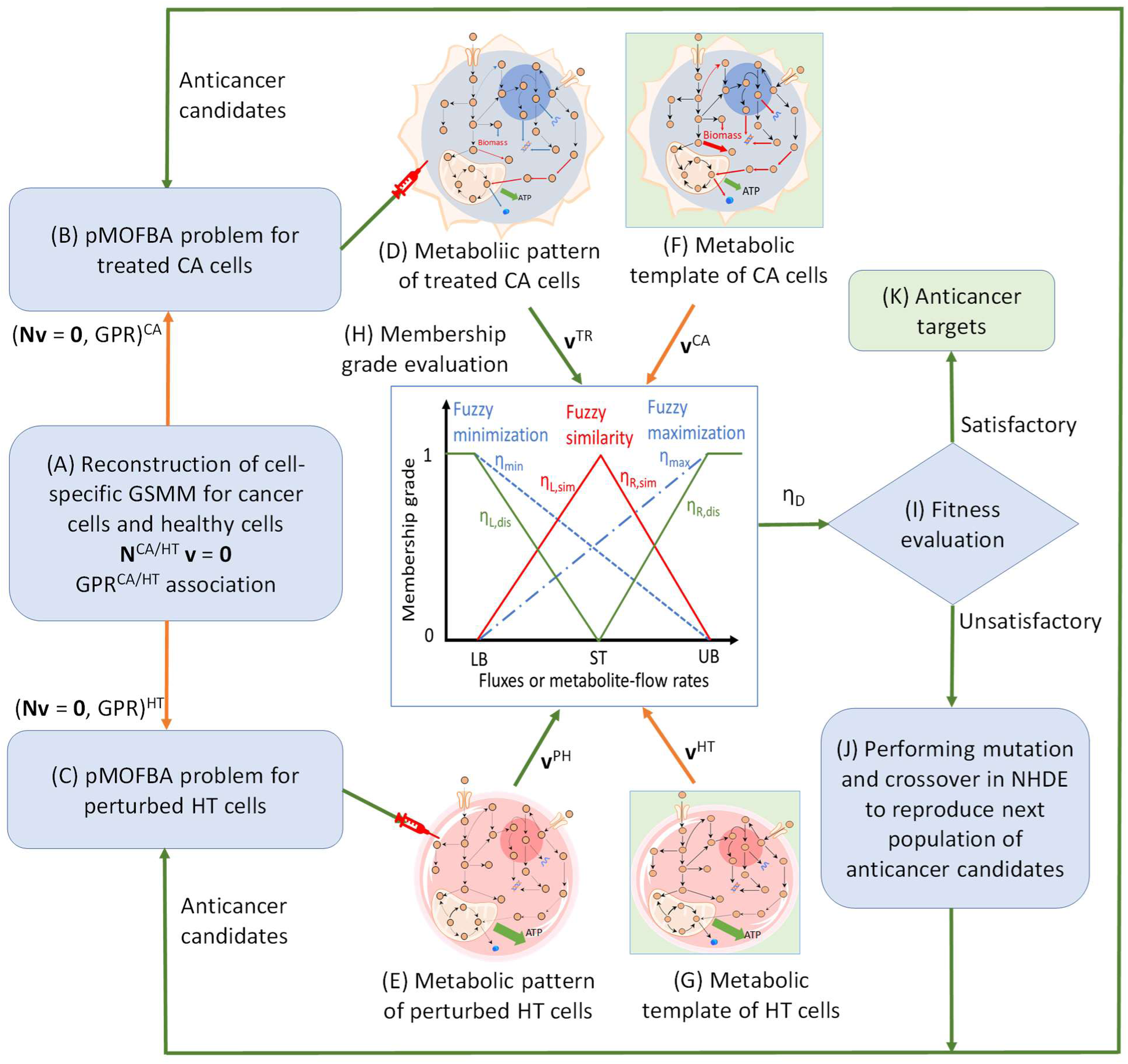
Constraint-based and fuzzy multi-objective optimization framework for anticancer target discovery. (A) Tissue-specific GSMMs are reconstructed for cancer-associated and corresponding healthy-tissue conditions. (B, C) Parsimonious multi-objective flux balance analysis is performed for each treated cancer and perturbed healthy-tissue model, respectively. (D, E) Metabolic patterns are calculated for each candidate perturbation. (F, G) Cancer-associated and healthy-tissue reference templates are obtained from clinical data when available or, otherwise, from unperturbed model simulations. (H) Flux and metabolite-flow values are transformed into a fuzzy decision score *η_D_*. (I) The fitness of each anticancer candidate is evaluated based on *η_D_* to guide target selection. (J) When the decision criterion is not satisfied, the nested hybrid differential evolution algorithm generates a new population of candidate perturbations. Steps B–J are repeated until the stopping criterion is met. (K) The highest-ranking candidate perturbations are retained as predicted anticancer targets.

#### 2.4.1. Evaluation of cell viability

The response of each treated cancer (TC) model was evaluated using the cancer-cell inhibition score, 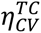, which integrates fuzzy membership grades representing the suppression of biomass production and ATP synthesis, as expressed in Equation (7).

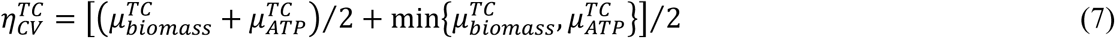

Here, the membership grades, 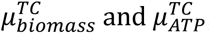, represent the extent of suppression of the biomass growth rate and ATP synthesis rate, respectively. Equation (7) combines their average performance with the smaller membership grade, thereby favouring treatments that simultaneously suppress cancer-cell growth and energy production rather than predominantly affecting only one objective. Although 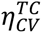 was originally termed the treated cancer cell-viability grade, a larger value indicates greater suppression of predicted cancer-cell viability and thus stronger treatment efficacy. Accordingly, this measure is referred to here as the cancer-cell inhibition score.

The response of each perturbed healthy-tissue (PH) model was assessed using the healthy-tissue viability-preservation score, 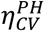 . For PH models, the objectives were to minimize unnecessary biomass production while preserving ATP synthesis, thereby emphasizing the maintenance of essential energetic and metabolic functions rather than proliferative activity. The formulation of 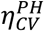 was analogous to Equation (7), but its membership functions were defined according to the healthy-tissue objectives. Consequently, a higher 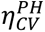 indicates better preservation of healthy-tissue metabolic function following target perturbation. The two scores, 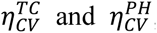, were incorporated into the ACTD framework to evaluate candidate enzyme targets in terms of predicted cancer-cell inhibition and healthy-tissue viability preservation, respectively.

#### 2.4.2. Metabolic-pattern comparison

More than 5,000 reaction fluxes and metabolite synthesis rates were calculated for the TC and PH models using the parsimonious multi-objective flux balance analysis formulation, with Ç_1_ = Ç_2_ = 1 in Equation (5). Collectively, these predicted fluxes and synthesis rates characterized the metabolic pattern of each model.

The metabolic patterns of TC and PH models were compared with those of untreated CA and HT reference models. Fuzzy similarity was used to quantify metabolic resemblance to the HT reference, whereas fuzzy dissimilarity measured the extent of separation from the CA reference. These membership grades were integrated into the metabolic-deviation score defined in Equation (8).

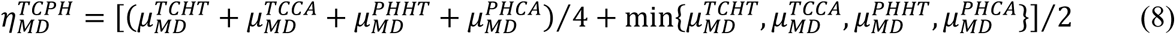

The membership grades, 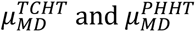, associated with the HT reference quantify the fuzzy similarity of TC and PH metabolic patterns to the healthy metabolic state. In contrast, the grades, 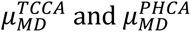, associated with the CA reference represent their fuzzy dissimilarity from the cancer-associated metabolic state. Equation (8) combines the average membership grade with the minimum grade to balance overall metabolic improvement against the least favourable response. A higher metabolic-deviation score therefore indicates a more favourable predicted treatment response, characterized by a shift of TC metabolism towards the healthy reference and away from the cancer-associated state, together with preservation of a healthy-like metabolic pattern in PH models.

Fuzzy similarity and dissimilarity were quantified using the two-sided membership functions defined in Equations (9) and (10), respectively.

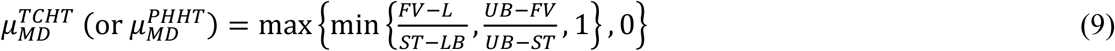

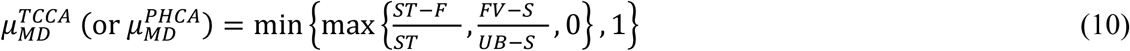

Here, *FV* denotes the calculated reaction flux or metabolite synthesis rate, whereas *LB*, *UB*, and *ST* represent the lower bound, upper bound, and reference value, respectively. These parameters were obtained from clinical measurements when available or otherwise derived from the CA and HT metabolic reference templates. In the present study, no direct clinical measurements were used to define the membership-function parameters. Instead, the reference values (*ST*) were derived from the corresponding CA and HT metabolic reference templates, with the lower and upper bounds assigned as *LB* = *ST*⁄4 and *UB* = 4*ST*, respectively. These parameters represent predefined modelling assumptions rather than experimentally calibrated physiological limits.

Finally, the fuzzy multi-objective hierarchical optimization problem illustrated in Figure 3A was transformed into a PBMFD formulation using the membership functions shown in Figure 3B. The resulting optimization problem was solved using the nested hybrid differential evolution (NHDE) algorithm [30, 34]. The theoretical conditions supporting this transformation have been described previously [34].

**Figure 3.**
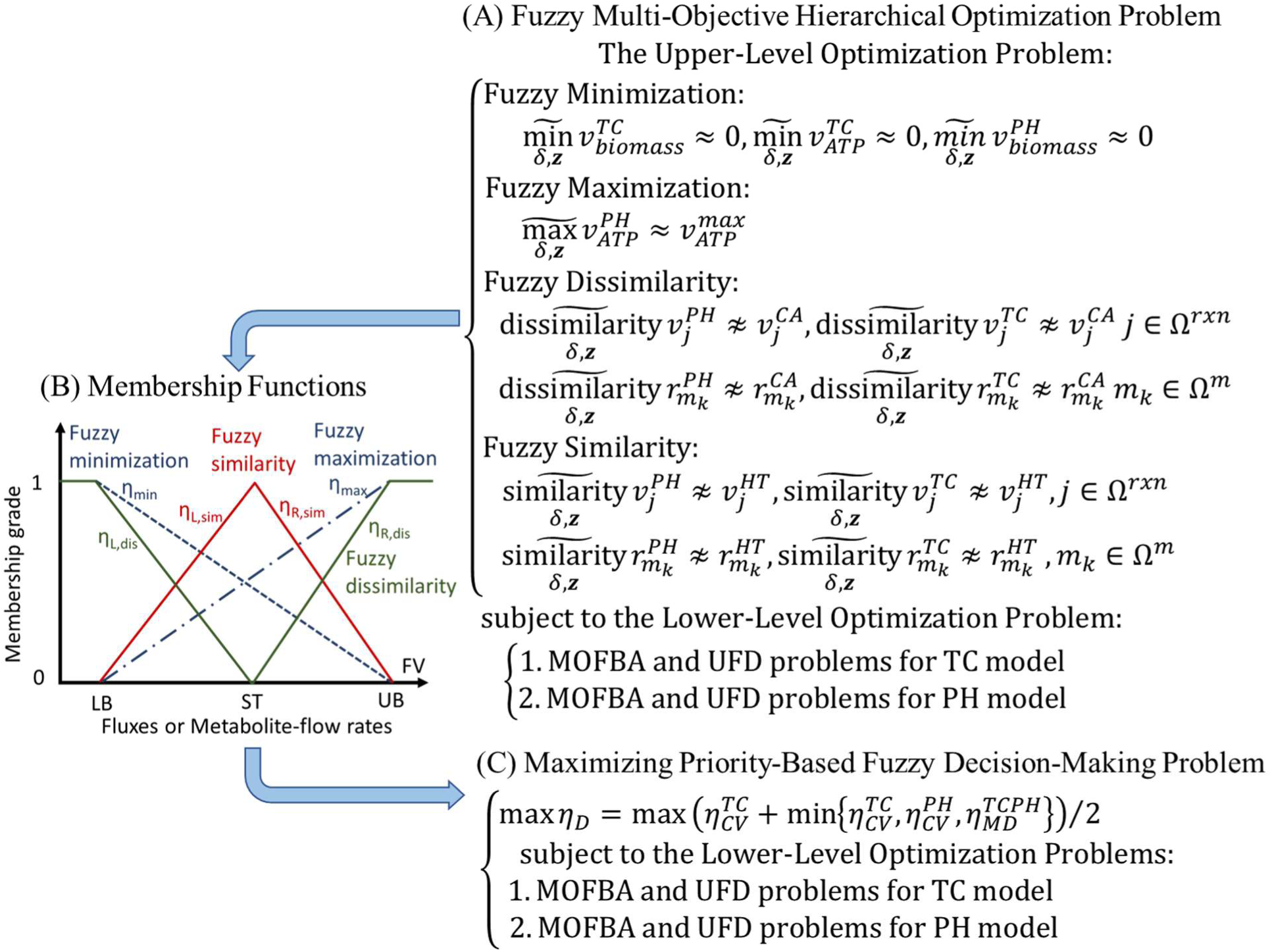
Transformation of the fuzzy multi-objective hierarchical optimization problem into a priority-based maximizing fuzzy decision problem. (A) Hierarchical optimization framework incorporating fuzzy minimization, maximization, similarity, and dissimilarity objectives for treated cancer-cell and perturbed healthy-tissue metabolic models. (B) Fuzzy membership functions used to quantify the degree of satisfaction of each objective. One-sided membership functions represent minimization and maximization objectives, whereas two-sided membership functions represent similarity and dissimilarity objectives. The reference value (ST) is obtained from clinical data when available or otherwise estimated from cancer-associated and healthy-tissue metabolic templates. The lower (LB) and upper (UB) bounds define the corresponding membership-function intervals and are set to LB = ST/4 and UB = 4ST in this study. (C) Transformation of the fuzzy multi-objective hierarchical formulation into a priority-based maximizing fuzzy decision problem by integrating individual membership grades according to their assigned priorities.

## 3. Results and discussion

### 3.1. Pan-cancer metabolic models

Cancer-associated (CA) and healthy-tissue (HT) genome-scale metabolic models were reconstructed for 33 cancer types using the CORDA algorithm [35]. The CA models retained approximately 62–69% of the metabolites and 55–60% of the reactions in Recon3D, whereas the reconstructed tissue-specific HT models retained approximately 62–77% and 53–67%, respectively. All models included nine subcellular compartments. Detailed reconstruction statistics are provided in Supplementary Table S1.

For nine cancer types lacking matched normal samples and skin cutaneous melanoma (SKCM), which contained only one normal sample, the generic Recon3D model was used as the HT reference. Comparisons involving these ten cancer types should therefore be interpreted cautiously because differences in network coverage and reference-model composition may contribute to the predicted metabolic alterations. Moreover, the numbers of retained metabolites and reactions characterize network structure rather than metabolic activity or predictive accuracy.

### 3.2. Metabolic alterations and robust candidate biomarkers

#### 3.2.1. Classification of differential metabolite-flow ranges

The metabolic models reconstructed for 33 cancer types were evaluated under five nutrient conditions, yielding 165 cancer–medium instances. Feasible ranges of intracellular metabolite synthesis, uptake, and secretion rates were determined using parsimonious multi-objective metabolite flow variability analysis (pMOMFVA). The predicted ranges for CA and HT models were compared and classified into seven categories according to their relative positions and degree of overlap: complete increase (CI), partial increase (PI), inclusive increase (II), unchanged (UC), inclusive decrease (ID), partial decrease (PD), and complete decrease (CD).

Positive classifications (CI, PI, and II) predominated in most cancer models, suggesting increased feasible metabolite-production capacity relative to the corresponding HT references. TGCT, BRCA, LAML, and THYM exhibited relatively large numbers of CI features, whereas PCPG, UCEC, LUSC, and LAML showed comparatively large numbers of CD features. Nevertheless, the distribution of classifications varied substantially among cancer types, reflecting heterogeneity in their predicted metabolic capabilities. Detailed classification counts are provided in Supplementary Table S2.

Among these categories, CI and CD provide the strongest model-based discrimination because the predicted CA and HT flux ranges do not overlap. However, these classifications represent differences in feasible metabolic ranges rather than experimentally measured metabolite distributions. Their interpretation also depends on model reconstruction, objective-function selection, nutrient constraints, and reference-model composition. Accordingly, the identified metabolic alterations are considered candidate biomarkers requiring further experimental and clinical validation.

#### 3.2.2. Shared candidate metabolic biomarkers across TCGA cancer types

Table 1 summarizes the classification counts and average scores for intracellular metabolite synthesis, secretion, and uptake across the 165 cancer–medium instances. The most prominent positive scores were associated with mitochondrial phospholipid metabolism, aromatic amino acid metabolism, and nutrient exchange.

**Table 1.** Classifications counts and average scores (Ave.) for intracellular metabolite synthesis, secretion, and uptake rates across 165 TCGA-derived cancer-medium model instances.

| Rates |  | CI (+3) | PI (+2) | II (+1) | UC (0) | ID (-1) | PD (-2) | CD (-3) | Ave. |
|---|---|---|---|---|---|---|---|---|---|
| Intracellular Metabolites | Cytidine-5'-Diphosphate-Diacylglycerol | 148 | 17 | 0 | 0 | 0 | 0 | 0 | 2.9 |
|  | L-Tryptophan | 136 | 28 | 0 | 1 | 0 | 0 | 0 | 2.81 |
|  | Cardiolipin | 134 | 26 | 0 | 5 | 0 | 0 | 0 | 2.75 |
|  | Phosphatidyl Glycerol Phosphate | 128 | 7 | 30 | 0 | 0 | 0 | 0 | 2.59 |
|  | 1-Phosphatidyl-1D-Myo-Inositol | 126 | 23 | 1 | 15 | 0 | 0 | 0 | 2.58 |
|  | L-Tyrosine | 113 | 45 | 0 | 0 | 7 | 0 | 0 | 2.56 |
|  | Myo-Inositol | 113 | 22 | 15 | 15 | 0 | 0 | 0 | 2.41 |
|  | Cytidine-5'-Monophosphate-N-Acetyl-Beta-Neuraminate | 79 | 79 | 4 | 1 | 0 | 0 | 2 | 2.38 |
|  | Phosphatidylglycerol | 109 | 4 | 52 | 0 | 0 | 0 | 0 | 2.35 |
|  | Deoxyguanosine-5'-Triphosphate | 41 | 123 | 0 | 0 | 0 | 1 | 0 | 2.22 |
|  | Diglyceride | 73 | 60 | 27 | 3 | 0 | 2 | 0 | 2.19 |
|  | Phosphatidylethanolamine | 53 | 93 | 14 | 1 | 1 | 3 | 0 | 2.13 |
|  | Cholesterol Sulfate | 84 | 60 | 5 | 0 | 2 | 13 | 1 | 2.1 |
|  | Phosphatidate | 68 | 66 | 11 | 20 | 0 | 0 | 0 | 2.1 |
|  | Oxidized Thioredoxin | 50 | 91 | 21 | 0 | 0 | 1 | 2 | 2.09 |
|  | Reduced Thioredoxin | 49 | 92 | 21 | 0 | 0 | 1 | 2 | 2.08 |
|  | Deoxyguanosine-5'-Diphosphate | 28 | 120 | 9 | 7 | 0 | 1 | 0 | 2.01 |
|  | Bicarbonate | 1 | 0 | 2 | 1 | 54 | 53 | 54 | -1.92 |
|  | Ubiquinol-10 | 0 | 0 | 3 | 4 | 46 | 62 | 50 | -1.92 |
|  | Ubiquinone-10 | 0 | 0 | 3 | 4 | 46 | 61 | 51 | -1.93 |
|  | 3'-S-Hydroxy-Pravastatin-CoA | 0 | 0 | 0 | 49 | 2 | 0 | 114 | -2.08 |
|  | 3'-S-Hydroxy-Pravastatin-Tetranor-CoA | 0 | 0 | 0 | 49 | 2 | 0 | 114 | -2.08 |
|  | 3'-S-Hydroxy-Pravastatin-Tetranor | 0 | 0 | 0 | 49 | 2 | 0 | 114 | -2.08 |
| Secretion | L-Lactate | 61 | 36 | 31 | 7 | 13 | 4 | 13 | 1.37 |
|  | 3-S-Hydroxy-Pravastatin-Tetranor | 0 | 0 | 0 | 49 | 2 | 0 | 114 | -2.08 |
| Uptake | L-Lysine | 157 | 8 | 0 | 0 | 0 | 0 | 0 | 2.95 |
|  | L-Tryptophan | 155 | 10 | 0 | 0 | 0 | 0 | 0 | 2.94 |
|  | L-Threonine | 127 | 36 | 2 | 0 | 0 | 0 | 0 | 2.76 |
|  | L-Isoleucine | 100 | 63 | 2 | 0 | 0 | 0 | 0 | 2.59 |
|  | L-Tyrosin | 116 | 41 | 1 | 0 | 7 | 0 | 0 | 2.57 |
|  | L-Methionine | 106 | 53 | 1 | 0 | 5 | 0 | 0 | 2.55 |
|  | Choline | 100 | 55 | 7 | 1 | 0 | 2 | 0 | 2.5 |
|  | L-Leucine | 85 | 77 | 2 | 0 | 1 | 0 | 0 | 2.48 |
|  | 3-Hydroxy Pravastatin | 0 | 0 | 0 | 49 | 2 | 0 | 114 | -2.08 |

Among intracellular metabolites, CDP-diacylglycerol (average score = 2.90), cardiolipin (2.75), and phosphatidylglycerol phosphate (2.59) exhibited consistently increased synthesis rates. CDP-diacylglycerol and phosphatidylglycerol phosphate are intermediates in cardiolipin biosynthesis, whereas cardiolipin is a major phospholipid of the mitochondrial inner membrane. Their coordinated increases suggest enhanced mitochondrial phospholipid synthesis or remodelling across the cancer models, potentially supporting mitochondrial membrane organization and bioenergetic adaptation. Nevertheless, increased synthetic flux does not necessarily indicate greater intracellular cardiolipin abundance, which also depends on degradation, acyl-chain remodelling, and mitochondrial turnover [36, 37].

L-Tryptophan (2.81) and L-tyrosine (2.56) also exhibited strong positive average scores. Together with their increased uptake rates, these findings suggest altered aromatic amino acid metabolism in the cancer models. These amino acids contribute to protein synthesis and serve as precursors for signalling and immunomodulatory metabolites. Because tryptophan is an essential amino acid and tyrosine can be produced through phenylalanine hydroxylation, the predicted intracellular synthesis variables should not be interpreted as evidence of unrestricted de novo amino acid biosynthesis. Instead, they may represent model-defined production, interconversion, or turnover fluxes.

In contrast, ubiquinone-10 and ubiquinol-10 both exhibited predominantly negative average scores (−1.92), indicating reduced feasible synthesis rates in CA models. However, these predictions do not establish global suppression of oxidative phosphorylation because cancer cells may retain substantial mitochondrial respiratory activity. Changes in ubiquinone-related synthesis may reflect pathway constraints, redox demands, compartmentalization, or cellular objective functions. Thus, the combination of increased cardiolipin-related synthesis and reduced ubiquinone-related synthesis is more appropriately interpreted as predicted remodelling of mitochondrial lipid and redox metabolism rather than uniform enhancement or suppression of electron transport [36].

Pravastatin-related metabolites also exhibited negative scores. Because these compounds are xenobiotics rather than endogenous human metabolites, their predicted synthesis and exchange rates depend on the inclusion of relevant reactions and substrates in Recon3D and the specified nutrient environments. These results should therefore be treated as xenobiotic-associated model predictions rather than evidence of altered statin pharmacokinetics or clinical drug clearance.

At the exchange-reaction level, lactate secretion increased in many cancer–medium instances, consistent with enhanced glycolytic carbon disposal and aerobic glycolysis [4–7]. However, its average score did not reach the threshold of 2.4 used to identify medium-robust candidate biomarkers. Lactate secretion should therefore be regarded as a recurrent pan-cancer trend rather than a strictly medium-robust biomarker.

Increased uptake rates were also predicted for lysine, tryptophan, threonine, isoleucine, tyrosine, methionine, and leucine across most cancer–medium instances. This pattern is consistent with the increased demand for amino acids associated with protein synthesis, biomass production, redox regulation, and growth-related signalling. Leucine and isoleucine may contribute to branched-chain amino acid metabolism and mTOR-associated anabolic signalling, whereas tryptophan uptake may support kynurenine-pathway metabolism. Increased uptake of several large neutral amino acids may involve LAT1/SLC7A5 or related transport systems, although transporter-specific mechanisms cannot be established from exchange-flux predictions alone [5, 38, 39].

Choline uptake was also elevated in most cancer–medium instances, suggesting increased demand for phosphatidylcholine synthesis through the Kennedy pathway. Phosphatidylcholine is a major component of cellular and organelle membranes, and increased choline availability may support membrane biogenesis, organelle expansion, and lipid-mediated signalling in proliferating cancer cells [40, 41]. However, experimental validation through choline-uptake measurements, transporter-expression analysis, choline kinase assays, isotope tracing, or lipidomics is required to establish the underlying mechanisms.

Collectively, these predictions suggest coordinated alterations in mitochondrial phospholipid metabolism, amino acid utilization, nutrient uptake, and glycolytic carbon disposal across multiple cancer types. Such changes are consistent with the anabolic demands and metabolic adaptations associated with malignant proliferation. Nevertheless, pMOMFVA predicts feasible steady-state metabolic fluxes rather than intracellular metabolite concentrations; therefore, the predicted alterations should not be interpreted as metabolite accumulation without independent experimental evidence.

The Cancer Cell Line Encyclopedia (CCLE) metabolomics dataset provides baseline metabolite profiles across human cancer cell lines and integrates metabolomic measurements with genomic alterations, gene expression, DNA methylation, and genetic dependency data [63]. A qualitative comparison of the predictions summarized in Table 1 with reported CCLE metabolite-abundance trends suggested concordant patterns for myo-inositol, lactate, lysine, tryptophan, threonine, isoleucine, tyrosine, methionine, choline, and leucine. Although this agreement provides preliminary support for the biological plausibility of the predicted alterations, metabolite abundance is not directly equivalent to synthesis, uptake, or secretion flux. Rigorous quantitative validation would require cell-line-specific metabolic models reconstructed using matched transcriptomic profiles and experimentally defined culture conditions.

#### 3.2.3. Sensitivity of metabolic alterations to objective functions and UFD reaction weighting in pMOMFVA

To assess the sensitivity of the predicted metabolic alterations to modelling assumptions, we evaluated three alternative configurations that varied the objective functions used in the CA models and the reaction-weighting scheme in the uniform flux distribution (UFD) formulation. The configurations and detailed classification results are provided in Supplementary Table 1 and Tables S2–S10.

The results indicated that the selection of CA-model objective functions influenced pMOMFVA classifications more strongly than the UFD reaction-weighting scheme. Compared with the reference formulation, which maximized both biomass and ATP production, maximizing biomass production alone substantially increased the number of CI classifications while reducing PI classifications (Tables S4 and S5). Nevertheless, several major metabolic alterations remained directionally consistent across the configurations, including increased cardiolipin-related synthesis and amino acid uptake, together with reduced ubiquinone-10 and ubiquinol-10 synthesis. These findings suggest that although individual classification outcomes depend on modeling assumptions, several prominent metabolic predictions remain robust.

#### 3.2.4. Cancer-type-associated and cancer-group-enriched candidate biomarkers

In addition to shared pan-cancer alterations, pMOMFVA identified metabolic features enriched in individual cancer types or restricted cancer groups (Table 2). Candidate biomarkers were screened by comparing the absolute average classification score for a selected cancer type, |*Ave*|_i_, with the corresponding absolute average score across the remaining cancer types. Features were retained when the selected cancer score was at least 2.4 and the comparison score was below 2.4. Because several metabolites were identified in more than one cancer type, these features are described as cancer-associated or cancer-group-enriched rather than strictly cancer-type-specific.

**Table 2.** Candidate metabolic biomarkers showing relative enrichment in individual cancer types or restricted groups of cancer types. Note: The threshold defines model-based relative enrichment rather than diagnostic specificity. Drug-related, dietary or plant-associated, and noncanonical metabolites should be curated and reported separately.

| Intracellular Metabolite Synthesis Rates | DMEM | HAM | HPLM | RPMI | VMH | Ave. | Cancer Type |
|---|---|---|---|---|---|---|---|
| Low Density Lipoprotein | 3 | 3 | 3 | 3 | 3 | 3 | BRCA |
| Low Density Lipoprotein | 3 | 3 | 3 | 3 | 3 | 3 | THCA |
| Cytidine-5'-DIphosphate | 2 | 3 | 2 | 3 | 2 | 2.4 | COAD |
| Formate | 2 | 3 | 2 | 2 | 3 | 2.4 | HNSC |
| Atorvastatin Acid | -3 | -3 | -3 | -3 | 0 | -2.4 | HNSC |
| Prostaglandin D2 | -3 | -3 | -3 | -3 | 0 | -2.4 | HNSC |
| Deoxythymidine-5'-Triphosphate | 3 | 3 | 3 | 3 | 3 | 3 | LIHC |
| Deoxycytidine | 3 | 3 | 3 | 3 | 3 | 3 | LIHC |
| Superoxide | -3 | -3 | -3 | -3 | 0 | -2.4 | LIHC |
| 7Alpha,12Alpha-Dihydroxy-5Beta-Cholestan-3-One | -3 | -3 | -3 | -3 | 0 | -2.4 | LIHC |
| 5Beta-Cholestane-3Alpha,7Alpha-Diol | -3 | -3 | -3 | -3 | 0 | -2.4 | LIHC |
| 5Beta-Cholestane-3Alpha,7Alpha,12Alpha-Triol | -3 | -3 | -3 | -3 | 0 | -2.4 | LIHC |
| 5Beta-Cholestane-3Alpha,7Alpha,26-Triol | -3 | -3 | -3 | -3 | -3 | -3 | LIHC |
| 3Alpha,7Alpha,12Alpha-Trihydroxy-5Beta-Cholestanoate | -3 | -3 | -3 | -3 | -3 | -3 | LIHC |
| L-Homoserine | 3 | 2 | 2 | 3 | 2 | 2.4 | LIHC |
| L-Homoserine | 3 | 3 | 3 | 3 | 3 | 3 | LUAD |
| Cytidine-5'-Triphosphate | 2 | 3 | 2 | 3 | 2 | 2.4 | PAAD |
| L-Cysteine | 3 | 2 | 3 | 2 | 2 | 2.4 | PAAD |
| (R)-3-Hydroxybutyrate | -3 | -3 | -3 | -1 | -3 | -2.6 | PRAD |
| (R)-3-Hydroxybutyrate | -3 | -3 | -3 | -1 | -3 | -2.6 | CESC |
| 1-Pyrroline-5-Carboxylate | -3 | -3 | -3 | -3 | 0 | -2.4 | CESC |
| Reduced Glutathione | -3 | -3 | -2 | -2 | -2 | -2.4 | THCA |
| Prostaglandin D2 | 0 | -3 | -3 | -3 | -3 | -2.4 | THCA |
| Prostaglandin E1 | 0 | -3 | -3 | -3 | -3 | -2.4 | THCA |
| (S)-Malate | -3 | -3 | -3 | -3 | -2 | -2.8 | STAD |
| (S)-Malate | -3 | -3 | -3 | -3 | -2 | -2.8 | LAML |
| (S)-Malate | -3 | -3 | -3 | -3 | -2 | -2.8 | LGG |
| Formaldehyde | 3 | 3 | 0 | 3 | 3 | 2.4 | LUSC |
| Glycine | 3 | 3 | 3 | 3 | 0 | 2.4 | DLBC |
| 2,4-Decadienoyl Coenzyme A | 3 | 3 | 3 | 3 | 0 | 2.4 | DLBC |
| 4-Decenoyl Coenzyme A | 3 | 3 | 3 | 3 | 0 | 2.4 | DLBC |
| Trans-4-Coumarate | 3 | 3 | 3 | 3 | 0 | 2.4 | OV |
| Trans-4-Coumarate | 3 | 3 | 3 | 3 | 0 | 2.4 | UVM |
| Hypoxanthine | -3 | -3 | -3 | 0 | -3 | -2.4 | CHOL |

Low-density-lipoprotein-associated flow rates were increased under all five nutrient conditions in BRCA and THCA. These predictions suggest altered cholesterol uptake, transport, or processing, potentially supporting membrane synthesis, organelle biogenesis, and lipid-raft-associated signalling. Cholesterol-derived 27-hydroxycholesterol has also been implicated in cancer proliferation and progression [42–44]. However, increased LDL-associated flow should not be interpreted directly as intracellular accumulation of intact LDL particles without examining the associated transport and cholesterol-processing reactions.

COAD exhibited increased CDP synthesis, suggesting alterations in cytidine-nucleotide metabolism. CDP participates in pyrimidine interconversion and contributes to phospholipid biosynthesis through CDP-activated intermediates. Increased CDP synthesis may therefore reflect altered nucleotide metabolism or membrane-lipid demand, although the present analysis cannot distinguish enhanced synthesis from reduced downstream utilization [5, 45].

In HNSC, increased formate synthesis was accompanied by reduced prostaglandin D2 and atorvastatin-associated flow. Increased formate is compatible with altered serine–glycine–one-carbon metabolism, which contributes to nucleotide biosynthesis, methylation reactions, and redox regulation [46, 47]. Reduced prostaglandin D2 synthesis suggests changes in eicosanoid metabolism, although its biological consequences depend on receptor expression and the immune microenvironment. By contrast, atorvastatin-associated flow represents a xenobiotic-related model prediction and should not be interpreted as an endogenous cancer biomarker.

LIHC exhibited increased dTTP and deoxycytidine synthesis rates, consistent with alterations in nucleotide metabolism associated with DNA synthesis and repair. Several cholesterol-derived bile-acid intermediates, including cholestane triols and 3alpha,7alpha,12alpha-trihydroxy-5beta-cholestanoate, showed reduced synthesis rates. These findings are compatible with diminished flux through differentiated hepatic bile-acid pathways relative to the healthy liver model, consistent with the involvement of bile-acid metabolism in liver-cancer biology [48, 49].

Reduced superoxide synthesis was also predicted in LIHC. However, a lower flux through superoxide-generating reactions does not necessarily imply reduced intracellular oxidative stress. Reactive oxygen species (ROS) levels depend on the balance between production and antioxidant removal, as well as the identity and subcellular localization of individual ROS species. Consequently, reduced superoxide-generating flux or ubiquinone-related synthesis cannot independently establish lower cellular ROS levels [50, 51].

Increased L-homoserine synthesis was predicted in LIHC and LUAD. This result requires particularly caution because L-homoserine is not a well-established intermediate of canonical human metabolism. Its biological interpretation therefore requires verification of the corresponding Recon3D reactions, metabolite compartments, reaction directionality, medium components, and GPR associations. Until these relationships are confirmed, L-homoserine should not be emphasized as a candidate clinical biomarker.

PAAD exhibited increased CTP and L-cysteine synthesis rates. Increased CTP synthesis is compatible with enhanced pyrimidine metabolism and demand for CDP-activated intermediates involved in membrane-lipid biosynthesis [52]. Increased cysteine synthesis may support protein production, coenzyme A metabolism, iron–sulfur cluster formation, and glutathione biosynthesis. However, this prediction does not necessarily establish increased antioxidant capacity, which also depends on glutathione turnover, NADPH availability, and antioxidant enzyme activity [53].

Reduced (R)-3-hydroxybutyrate synthesis in PRAD and CESC suggests altered ketone-body metabolism, whereas reduced 1-pyrroline-5-carboxylate synthesis in CESC may reflect changes in proline metabolism and redox coupling [54]. THCA exhibited reduced synthesis rates of glutathione, prostaglandin D2, and prostaglandin E1, suggesting alterations in redox regulation and eicosanoid metabolism [55]. Nevertheless, reduced glutathione synthesis cannot distinguish decreased production from altered oxidation, recycling, or turnover.

Reduced (S)-malate synthesis in STAD, LAML, and LGG may reflect changes in tricarboxylic acid cycle activity, the malate–aspartate shuttle, or cytosolic redox metabolism. Reduced hypoxanthine synthesis in CHOL suggests altered purine salvage or downstream utilization, whereas increased glycine synthesis in DLBC is compatible with enhanced one-carbon metabolism and nucleotide biosynthesis [56].

Several additional predictions, including increased formaldehyde, trans-4-coumarate, and unusual unsaturated acyl-CoA species, require further biochemical annotation. Formaldehyde may reflect altered one-carbon metabolism or aldehyde detoxification, but its biological effects are strongly influenced by detoxification capacity. Trans-4-coumarate is primarily associated with plant phenylpropanoid metabolism and should not be proposed as a human cancer biomarker without confirming the corresponding reactions and substrate sources. Similarly, the biological interpretation of 2,4-decadienoyl-CoA and 4-decenoyl-CoA requires verification of their reaction identifiers and pathway assignments.

Overall, the cancer-associated features summarized in Table 2 suggest heterogeneous alterations in nucleotide, lipid, one-carbon, redox, eicosanoid, ketone-body, and purine metabolism. Although several predictions involve well-established endogenous human metabolites, others depend on xenobiotic-associated or incompletely characterized reactions. Reaction-level annotation and medium-composition verification are therefore essential before prioritizing these features for experimental biomarker validation.

### 3.3. Identification of predicted anticancer targets

The ACTD framework was applied to 33 cancer types under five nutrient conditions to identify candidate enzyme perturbations with predicted anticancer activity. Candidate targets were evaluated using the priority-based fitness function illustrated in Figure 3C, which incorporated three scores ranging from 0 to 1: cancer-cell inhibition 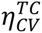, healthy-tissue viability preservation 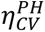, and metabolic deviation 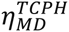.

Higher cancer-cell inhibition scores indicated stronger predicted suppression of biomass production and ATP synthesis in treated cancer (TC) models. Higher healthy-tissue viability-preservation scores represented better maintenance of the specified metabolic objectives in perturbed healthy-tissue (PH) models. Higher metabolic-deviation scores indicated a more favourable predicted metabolic response, characterized by greater separation of TC metabolism from the untreated CA reference and preservation of healthy-like metabolic patterns in PH models Although the cancer-cell inhibition score was originally termed a cell-viability grade, higher values correspond to lower predicted cancer-cell viability and therefore greater treatment efficacy. The healthy-tissue viability-preservation score remained close to unity for the optimal candidate targets and provided limited discrimination among them; accordingly, it is not presented in the following tables. The metabolic-deviation score provides complementary information about the broader metabolic consequences of each intervention. However, these model-derived scores should not be interpreted as direct experimental measures of treatment efficacy, therapeutic selectivity, or toxicity.

#### 3.3.1. Predicted broad-coverage enzyme targets

A total of 56 feasible single-enzyme perturbations satisfied the computational screening requirements in at least one cancer-medium instance. Table 3 summarizes the 18 highest-ranking targets according to their treatment-success counts, with treatment success defined by a cancer-cell inhibition score of 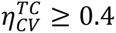.

**Table 3.** Treatment-Success Counts of Enzyme Targets Identified from 165 Cancer-Specific Metabolic Models. Note: Treatment success is defined as 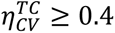.

| Gene Symbol | Encoded Enzyme | Superpathway | Pathway | Success Count |
|---|---|---|---|---|
| CRLS1 | Cardiolipin synthase 1 | Lipid metabolism | Glycerophospholipid metabolism | 165 |
| PGS1 | Phosphatidylglycerophosphate synthase 1 | Lipid metabolism | Glycerophospholipid metabolism | 163 |
| PTPMT1 | Protein tyrosine phosphatase mitochondrial 1 | Lipid metabolism | Glycerophospholipid metabolism | 148 |
| ADSL | Adenylosuccinate lyase | Nucleotide metabolism | Purine metabolism | 122 |
| ADSS2 | Adenylosuccinate synthase 2 | Nucleotide metabolism | Purine metabolism | 122 |
| DHODH | Dihydroorotate dehydrogenase (quinone) | Nucleotide metabolism | Pyrimidine metabolism | 117 |
| GK | Glycerol kinase | Lipid metabolism | Glycerolipid metabolism | 86 |
| PMVK | Phosphomevalonate kinase | Lipid metabolism | Sterol/terpenoid backbone biosynthesis | 72 |
| MVD | Mevalonate diphosphate decarboxylase | Lipid metabolism | Sterol/terpenoid backbone biosynthesis | 71 |
| MVK | Mevalonate kinase | Lipid metabolism | Sterol/terpenoid backbone biosynthesis | 71 |
| TYMS | Thymidylate synthetase | Nucleotide metabolism | Pyrimidine metabolism | 65 |
| LSS | Lanosterol synthase | Lipid metabolism | Steroid biosynthesis | 53 |
| SC5D | Sterol-C5-desaturase | Lipid metabolism | Steroid biosynthesis | 53 |
| SQLE | Squalene epoxidase | Lipid metabolism | Steroid biosynthesis | 53 |
| EBP | EBP cholestenol delta-isomerase | Lipid metabolism | Steroid biosynthesis | 51 |
| NSDHL | NAD(P) dependent steroid dehydrogenase-like | Lipid metabolism | Steroid biosynthesis | 35 |
| CEPT1 | Choline/ethanolamine phosphotransferase 1 | Lipid metabolism | Glycerophospholipid metabolism | 30 |
| CTPS1 | CTP synthase 1 | Nucleotide metabolism | Pyrimidine metabolism | 30 |

Among these targets, CRLS1, PGS1, and PTPMT1 exhibited the broadest predicted treatment coverage. All three enzymes participate in mitochondrial glycerophospholipid metabolism and cardiolipin biosynthesis. CRLS1 inhibition satisfied the predefined treatment-success criterion in all 165 cancer–medium instances, whereas PGS1 and PTPMT1 achieved success in 163 and 148 instances, respectively. These findings identify mitochondrial cardiolipin biosynthesis as a recurrent metabolic vulnerability across the reconstructed cancer models.

CRLS1 catalyses the terminal step of cardiolipin biosynthesis, whereas PGS1 and PTPMT1 participate in the synthesis and processing of phosphatidylglycerol phosphate [36, 37]. Perturbation of these enzymes may impair mitochondrial inner-membrane organization, respiratory-chain stability, oxidative phosphorylation, ATP production, and redox homeostasis. Their broad predicted effectiveness is therefore consistent with the importance of cardiolipin metabolism in mitochondrial function.

Figure 4 presents the cancer-cell inhibition and metabolic-deviation scores following CRLS1 inhibition across 33 cancer types and five nutrient conditions. Strong predicted inhibition was observed in most cancer types, with minimum inhibition scores approaching unity in a substantial proportion of the models. UVM, KIRC, BRCA, and HNSC exhibited consistently high inhibition scores across all five media. By contrast, MESO, SARC, and PCPG showed the lowest minimum scores, approximately 0.50, 0.55, and 0.59, respectively, whereas DLBC had a minimum score of approximately 0.71. MESO and SARC also exhibited substantial nutrient-dependent variability, with maximum scores approaching unity, while PCPG showed a lower maximum score of approximately 0.75. Nevertheless, all 165 cancer–medium instances satisfied the predefined treatment-success criterion, supporting the broad computational effectiveness of CRLS1 inhibition under the modelled conditions.

**Figure 4.**
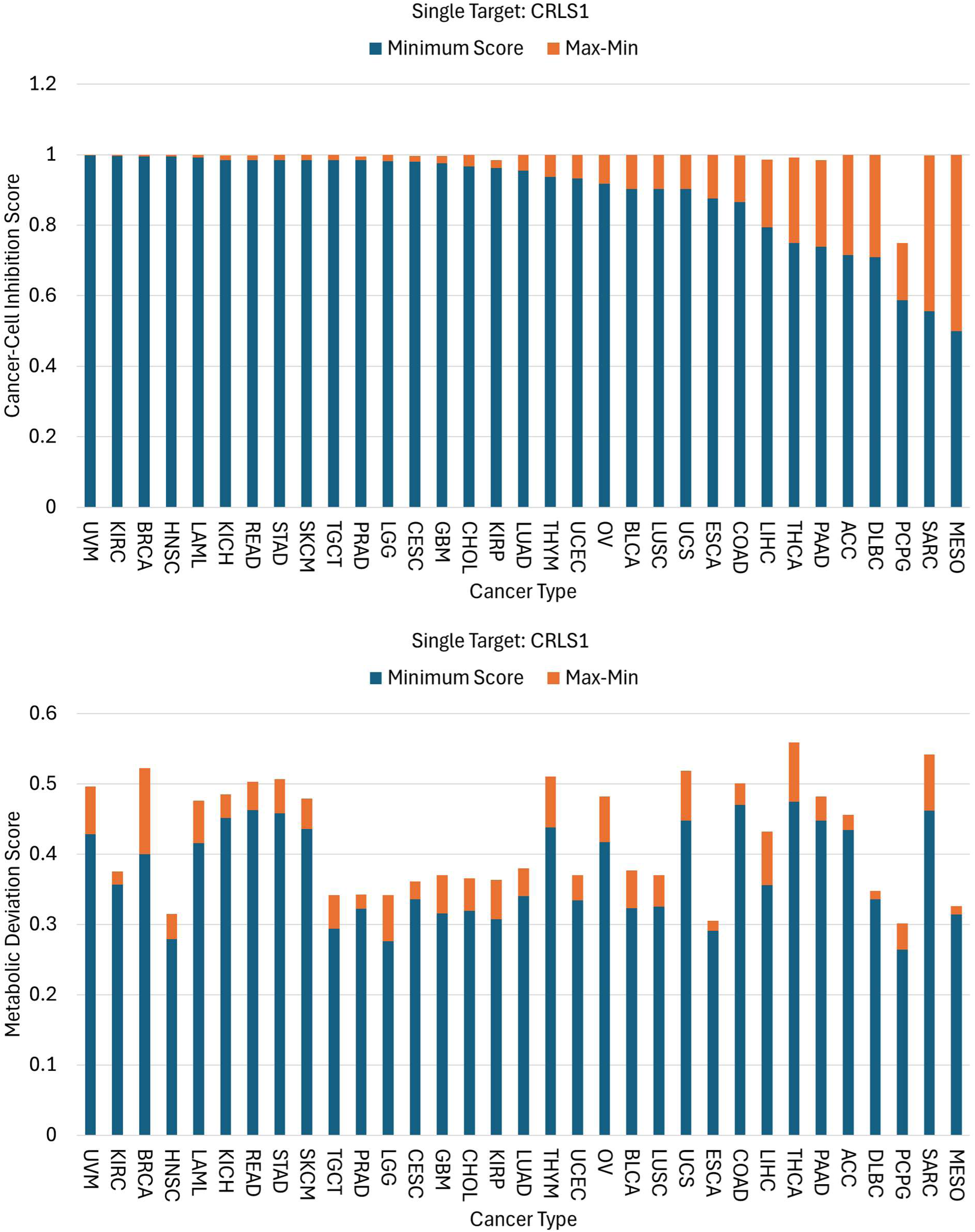
Predicted cancer-cell inhibition and metabolic-deviation scores following CRLS1 inhibition across 33 cancer types and five nutrient media. Blue bars represent the minimum score across the five media, and orange segments indicate the range from the minimum to the maximum score. Higher 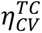 values indicate stronger predicted suppression of cancer-cell biomass and ATP production. Higher 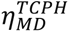 values indicate a more favourable metabolic response relative to the CA and HT reference templates.

To assess the sensitivity of this prediction to modelling assumptions, three alternative configurations (Case 1-3) were compared with the reference formulation (Case 0) across all cancer types and nutrient conditions (Figure 4; Supplementary Figures S1–S3). Case 1 retained the biomass- and ATP-production objectives but employed uniform UFD reaction weights, whereas Cases 2 and 3 used biomass-only objectives with expression-dependent and uniform UFD reaction weights, respectively. Under the bi-objective formulations (Figure 4; Supplementary Figure S1), CRLS1 inhibition produced high inhibition scores across most cancer types, although their magnitudes and nutrient-dependent variability differed. In contrast, the biomass-only formulations (Supplementary Figure S2 and S3) yielded inhibition scores approaching unity for nearly all cancer types, with PCPG remaining a notable exception. Differences between UFD weighting schemes were generally smaller than those associated with objective-function selection. Metabolic-deviation scores exhibited greater heterogeneity across cancer types and nutrient conditions, with broadly similar patterns among the four configurations.

These results indicate that the predicted inhibitory efficacy of CRLS1 is more sensitive to the selection of CA-model objective functions than to the UFD reaction-weighting scheme. Although CRLS1 retained broad predicted treatment coverage, the magnitude and nutrient dependence of its inhibitory effects varied with the objective-function formulation.

To assess the biological relevance of the predicted targets, genetic-dependency data from DepMap were analysed across ten cancer types. Among 582 cancer cell lines, dependencies on CRLS1, PGS1, and PTPMT1 were identified in 296 (50.9%), 350 (60.1%), and 294 (50.5%) cell lines, respectively (Supplementary Table S11). Notably, PGS1 exhibited a higher frequency of genetic dependency than CRLS1 in nine of the ten cancer types, despite the broader predicted efficacy of CRLS1 in the constraint-based simulations. This discrepancy highlights the distinction between experimentally observed genetic dependencies and model-predicted metabolic vulnerabilities, potentially reflecting differences in cellular context, network constraints, and nutrient availability. Nevertheless, the DepMap findings provide independent support for the biological relevance of mitochondrial cardiolipin biosynthesis as a potential anticancer target pathway.

Although PGS1 exhibited broad predicted coverage, it failed to satisfy the predefined inhibition criterion of 0.4 for SARC under HAM and RPMI media, while meeting the less stringent threshold of 0.3. PTPMT1 failed to meet the criterion in several cancer types, including DLBC, LGG, and TGCT. These differences may arise from cancer-specific metabolic network structures, nutrient-dependent constraints, or alternative phosphatidylglycerol-associated pathways.

Other targets, including ADSL, ADSS2, DHODH, GK, PMVK, MVD, MVK, TYMS, LSS, SC5D, SQLE, EBP, NSDHL, CEPT1, and CTPS1, exhibited lower treatment coverage, suggesting greater dependence on cancer type and nutrient availability. Functionally, ADSL and ADSS2 participate in purine metabolism, whereas DHODH, TYMS, and CTPS1 contribute to pyrimidine metabolism. GK, CEPT1, and enzymes involved in mevalonate and sterol biosynthesis participate in lipid metabolism. These findings identify nucleotide biosynthesis, glycerophospholipid metabolism, and sterol metabolism as additional candidate vulnerabilities, although their predicted effects are less consistent across cancer types and nutrient conditions.

Collectively, the computational and genetic-dependency analyses support mitochondrial cardiolipin biosynthesis as a promising pathway for further anticancer investigation. CRLS1 emerged as the most broadly effective predicted single-enzyme target, although its apparent efficacy depended partly on modelling assumptions. Because cardiolipin biosynthesis is also essential for normal mitochondrial function, the therapeutic selectivity of CRLS1 inhibition remains uncertain. Further experimental studies using cancer and nonmalignant cellular models are required to evaluate its anticancer efficacy and potential adverse metabolic effects.

#### 3.3.2. Predicted dual-perturbation combinations

The ACTD framework was extended to identify CRLS1-based combination interventions involving metabolite-associated reaction constraints and nutrient-uptake perturbations, with the aim of improving both cancer-cell inhibition and metabolic-deviation scores. Figure 5 summarizes the predicted responses to selected dual combinations involving CRLS1 inhibition and constraints associated with deoxycytidine 5’-diphosphate (dCDP), deoxythymidine 5’-diphosphate (dTDP), isopentenyl diphosphate (IPDP), or nicotinamide adenine dinucleotide phosphate (NADPH).

**Figure 5.**
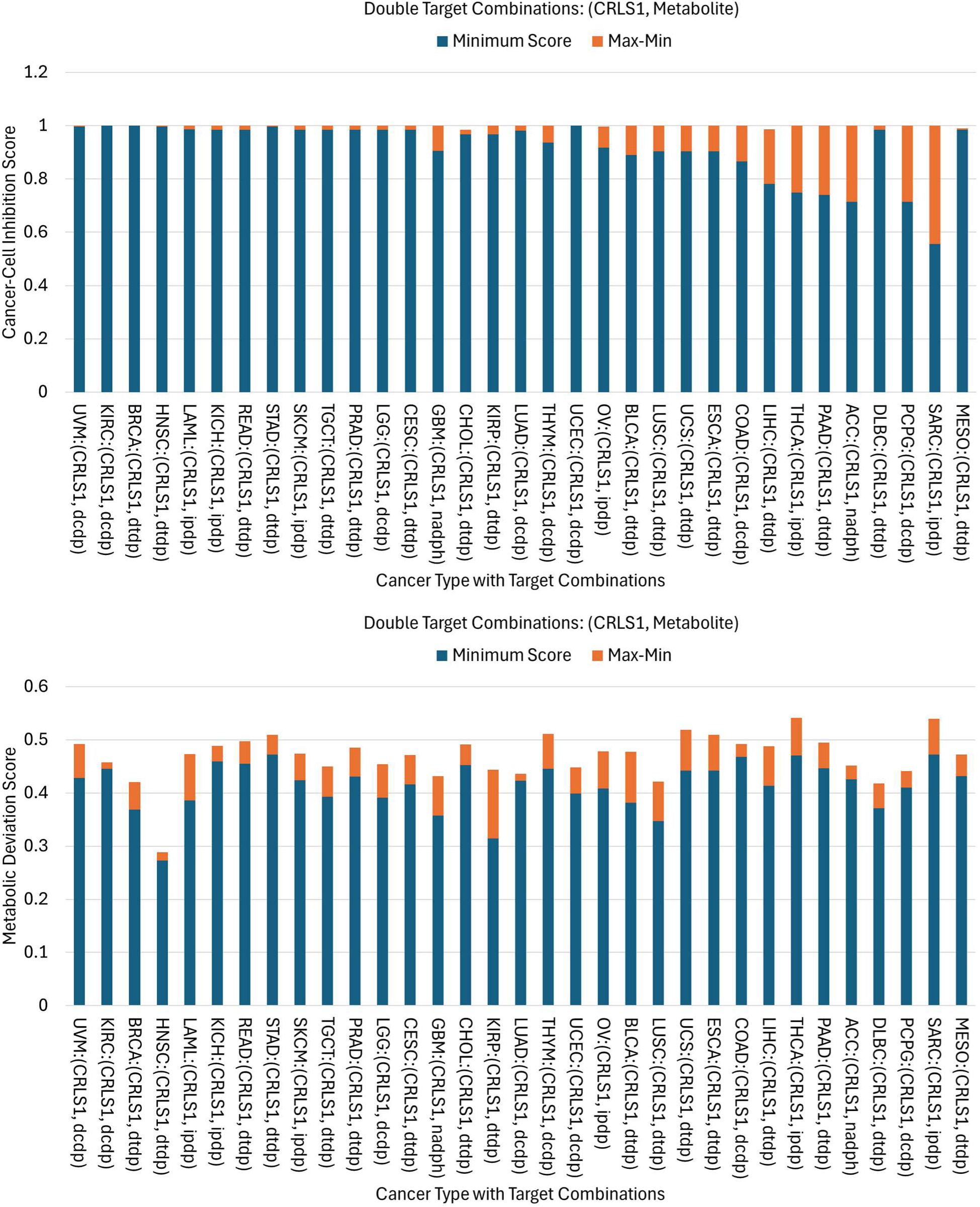
Predicted cancer-cell inhibition and metabolic-deviation scores for CRLS1-based dual perturbations across 33 cancer types and five nutrient media. Dual perturbations combined CRLS1 inhibition with model constraints involving deoxycytidine5′-diphosphate (dCDP), deoxythymidine 5′-diphosphate (dTDP), isopentenyl diphosphate (IPDP) or NADPH. Blue bars represent the minimum score across the five media, and orange segments indicate the range from the minimum to the maximum score.

The selected dual combinations produced high cancer-cell inhibition scores across most cancer types, with minimum values approaching unity under the five nutrient conditions. UVM, KIRC, BRCA, and HNSC exhibited consistently strong predicted inhibition. Compared with CRLS1 inhibition alone, the combinations also improved responses in MESO and DLBC, where minimum inhibition scores approached unity. Nevertheless, treatment efficacy remained heterogeneous. SARC exhibited the lowest minimum inhibition score (approximately 0.55), followed by PCPG and ACC (approximately 0.71), PAAD (0.74), and THCA (0.75). These results suggest that combining CRLS1 inhibition with additional metabolic perturbations can enhance predicted anticancer efficacy in selected cancer types, although responses remain sensitive to cancer-specific metabolic characteristics and nutrient conditions.

The enhanced effects of these combinations are biologically plausible at the metabolic-network level. CRLS1 inhibition may restrict cardiolipin biosynthesis and impair mitochondrial membrane organization and energy metabolism. Additional perturbations involving dCDP- or dTDP-associated reactions may disrupt nucleotide metabolism and precursor availability for DNA synthesis. Similarly, IPDP-associated constraints may influence isoprenoid and sterol biosynthesis, whereas NADPH-associated perturbations may affect reductive biosynthesis and cellular antioxidant capacity. Thus, simultaneous interference with mitochondrial phospholipid metabolism and complementary biosynthetic pathways may strengthen cancer-cell suppression.

Nevertheless, metabolite-associated perturbations should not be interpreted as direct pharmacological inhibition of the corresponding metabolites. Their effects depend on the implementation of model constraints, including changes in synthesis, consumption, or allowable reaction fluxes. In particular, NADPH participates in numerous metabolic pathways, and a broad NADPH-associated constraint may produce more extensive effects than inhibition of an individual enzyme [45]. Therefore, these computational interventions require further mapping to experimentally actionable enzymes or transporters.

Despite generally strong cancer-cell inhibition, metabolic-deviation scores remained heterogeneous, ranging approximately from 0.28 to 0.54 across cancer types and nutrient conditions (Figure 5). Several combinations achieved nearly complete predicted cancer-cell inhibition but only moderate metabolic-deviation scores, indicating that strong suppression of growth and ATP production did not necessarily correspond to equally favourable changes in the overall metabolic state. Furthermore, variation across nutrient conditions suggests that treatment-induced metabolic responses depend partly on nutrient availability.

These findings emphasize the importance of simultaneously evaluating cancer-cell inhibition and metabolic deviation when prioritizing combination interventions. Improvements in predicted anticancer efficacy alone are insufficient to establish favourable overall metabolic responses or therapeutic selectivity.

#### 3.3.3. Predicted triple-perturbation combinations

The ACTD framework was further extended to triple combinations by incorporating nutrient-uptake perturbations involving 3-phospho-D-glycerate (3PG), deoxyinosine (dIno), galactitol, D-glyceraldehyde, D-glycerate, hexanoate, L-kynurenine, or L-xylulose. Figure 6 summarizes the predicted cancer-cell inhibition and metabolic-deviation scores across 33 cancer types and five nutrient conditions.

**Figure 6.**
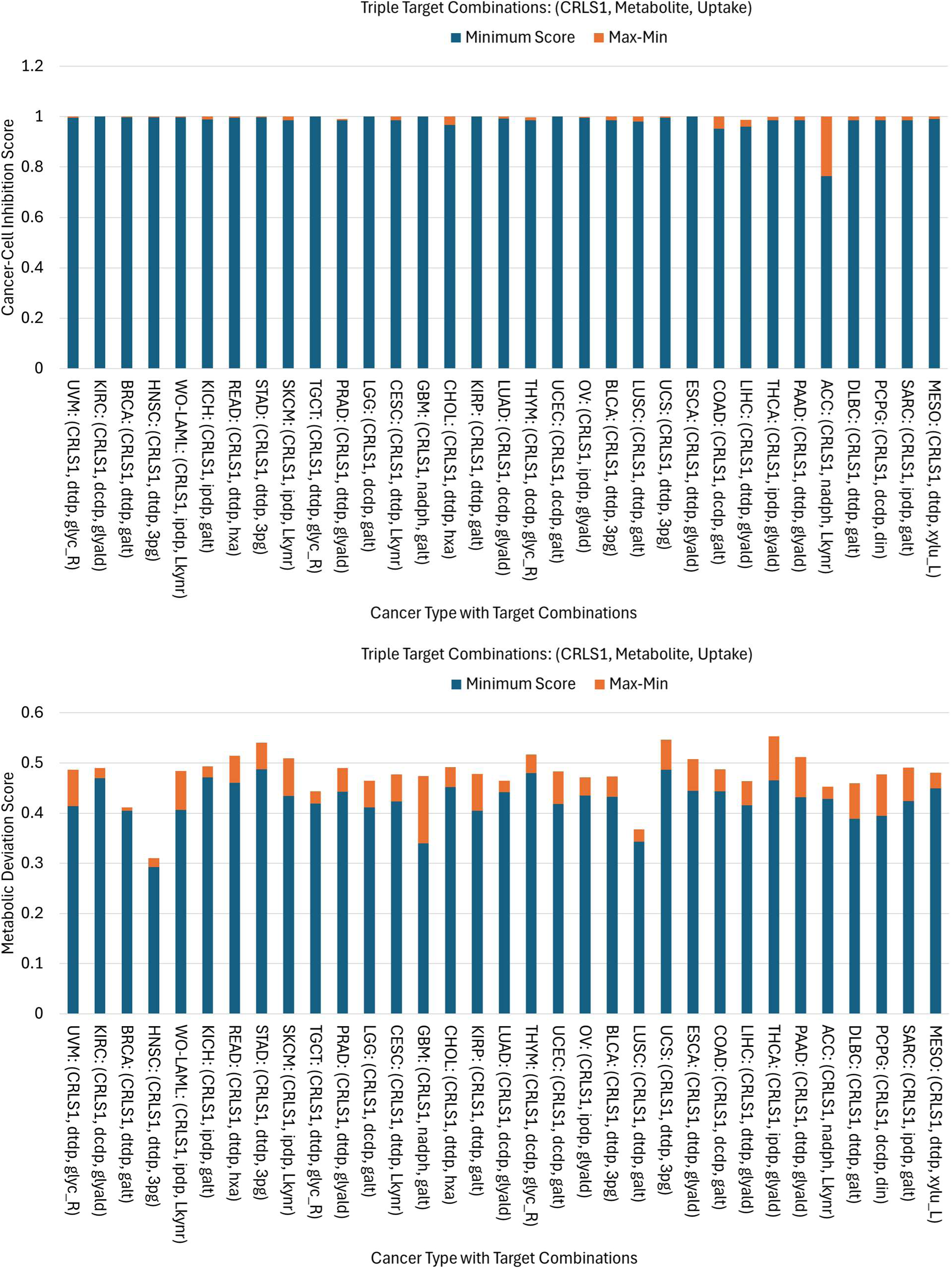
Predicted cancer-cell inhibition and metabolic-deviation scores for CRLS1-based triple perturbations across 33 cancer types and five nutrient media. Triple perturbations combined CRLS1 inhibition, a metabolite-associated constraint and an uptake perturbation. Uptake perturbations involved 3-phospho-D-glycerate (3pg), deoxyinosine (din), galactitol (galt), D-glyceraldehyde (glyald), D-glycerate (glyc_R), hexanoate (hxa), L-kynurenine (L-kymr) or L-xylulose (xylu_L). Blue bars represent the minimum score across the five media, and orange segments indicate the range from the minimum to the maximum score.

Compared with the selected dual combinations, the triple combinations generally exhibited more consistent inhibitory effects, with minimum cancer-cell inhibition scores approaching 0.97–1.0 for most cancer types. Notably, SARC and PCPG, which showed relatively weak or nutrient-dependent responses to dual perturbations, exhibited nearly complete predicted inhibition under their respective triple combinations. These findings suggest that additional nutrient-uptake perturbations may enhance the computational effectiveness of CRLS1-based interventions. However, because the metabolite-associated constraints may also differ between the selected dual and triple combinations, the improvements cannot necessarily be attributed to nutrient-uptake perturbations alone.

Despite these improvements, ACC exhibited incomplete inhibition under the combination comprising CRLS1 inhibition, an NADPH-associated perturbation, and increased L-kynurenine uptake. The predicted inhibition scores were approximately 0.780, 0.784, and 0.764 under DMEM, HAM, and RPMI media, respectively. In contrast, an alternative ACC-specific combination involving CRLS1 inhibition, glycerol kinase (GK) inhibition, and increased L-xylulose uptake achieved higher inhibition scores of approximately 0.985–1.0 across all five media. This result suggests that alternative combinations targeting complementary metabolic pathways may overcome limited responses in particular cancer–medium instances.

The enhanced effects of the CRLS1/GK/L-xylulose combination may arise from complementary perturbations of glycerolipid metabolism and mitochondrial cardiolipin biosynthesis. GK catalyses the phosphorylation of glycerol to glycerol 3-phosphate, an important precursor for glycerolipid and glycerophospholipid synthesis [58–60]. Combined inhibition of GK and CRLS1 may therefore restrict both glycerolipid precursor formation and cardiolipin biosynthesis, potentially disrupting cellular membrane-lipid metabolism and mitochondrial function.

Increased L-xylulose uptake may additionally modify carbon flux through glucuronate- and pentose-related pathways. However, its contribution to the predicted treatment response remains uncertain and requires further examination of the associated reactions and carbon-flow redistribution. The experimental feasibility of enhancing L-xylulose uptake and reproducing the predicted metabolic effects also remains to be established.

Although triple combinations generally achieved higher and more consistent cancer-cell inhibition scores, their metabolic-deviation scores remained variable, ranging approximately from 0.30 to 0.56 across cancer types and nutrient conditions (Figure 6). The relatively modest changes in metabolic-deviation scores, despite substantial improvements in inhibition for selected cancer types, indicate that stronger suppression of cancer-cell growth does not necessarily produce a more favourable overall metabolic response. Thus, cancer-cell inhibition and metabolic deviation should be evaluated jointly when prioritizing candidate combination strategies.

Overall, the combination analysis suggests that simultaneous perturbation of mitochondrial cardiolipin biosynthesis and nucleotide-, redox-, glycerolipid-, or nutrient-uptake-associated pathways can enhance predicted cancer-cell inhibition. The selected triple combinations generally outperformed the dual combinations in terms of inhibitory consistency, particularly in cancer types with incomplete responses to dual interventions. Nevertheless, heterogeneous metabolic-deviation scores highlight the potential trade-offs between cancer-cell suppression and broader metabolic responses. Moreover, metabolite-associated and nutrient-uptake perturbations represent computational interventions rather than directly actionable therapeutic strategies. Their translation into anticancer treatments requires identification of specific enzymes, transporters, or delivery mechanisms capable of reproducing the predicted effects, followed by experimental validation of therapeutic efficacy and selectivity.

## 4. Conclusions

This study developed an integrated constraint-based framework for pan-cancer metabolic biomarker discovery and anticancer target identification. Thirty-three TCGA-derived cancer models were evaluated under five nutrient conditions, yielding 165 cancer–medium instances. The pMOMFVA framework identified candidate metabolic alterations that were consistent across nutrient environments. Prominent predictions included increased synthesis rates of CDP-diacylglycerol, cardiolipin, phosphatidylglycerol phosphate, L-tryptophan, and L-tyrosine, together with enhanced uptake of choline and essential amino acids. These alterations suggest increased anabolic demand, membrane biosynthesis, and mitochondrial phospholipid remodelling. In contrast, reduced ubiquinone-10 and ubiquinol-10 synthesis rates indicated alterations in mitochondrial redox metabolism. Pravastatin-associated changes were also identified but should be regarded as xenobiotic-related model predictions rather than endogenous metabolic biomarkers.

Cancer-associated and cancer-group-enriched predictions further revealed metabolic heterogeneity among tumour types, including increased LDL-associated flux in breast and thyroid cancers, altered cytidine-nucleotide metabolism in colon and liver cancers, and reduced bile-acid- or tricarboxylic-acid-cycle-associated fluxes in selected cancers. Because the models predict feasible metabolite synthesis and exchange rates rather than intracellular concentrations, these alterations should be considered candidate biomarkers requiring experimental validation.

The ACTD framework further demonstrated the potential of fuzzy multi-objective optimization to prioritize enzyme perturbations by balancing predicted cancer-cell inhibition, ATP suppression, preservation of healthy-tissue metabolism, and robustness across nutrient conditions. Among 56 feasible single-enzyme perturbations, CRLS1 exhibited the broadest predicted anticancer effectiveness, satisfying the predefined treatment criterion in all 165 cancer–medium instances. PGS1 and PTPMT1 also showed broad treatment coverage, highlighting mitochondrial cardiolipin biosynthesis as a potential shared metabolic vulnerability. Genetic-dependency data from DepMap provided additional support for the biological relevance of these enzymes, although their experimental dependency patterns differed from the model-predicted treatment rankings. Furthermore, selected dual and triple perturbations improved predicted cancer-cell inhibition in several cancer types, but their metabolic-deviation scores remained heterogeneous, indicating that stronger inhibition did not necessarily correspond to more favourable overall metabolic responses.

Sensitivity analysis across four model configurations revealed that the predicted inhibitory efficacy of CRLS1 was more strongly influenced by cancer-cell objective-function selection than by UFD reaction weighting. Biomass-only formulations generally yielded near-maximal inhibition scores, whereas combined biomass- and ATP-production objectives revealed greater variability across cancer types and nutrient conditions. These findings support the broad predicted effectiveness of CRLS1 while emphasizing the importance of objective-function formulation when assessing treatment efficacy and nutrient-dependent robustness.

Overall, integrating pMOMFVA-based biomarker analysis with ACTD-based fuzzy optimization provides a mechanistically interpretable computational framework for identifying robust metabolic alterations and broad-coverage candidate anticancer targets. The findings highlight mitochondrial phospholipid metabolism, amino acid acquisition, nucleotide metabolism, and redox regulation as important pathways for further investigation. Future studies should validate these predictions through metabolomics, isotope tracing, genetic perturbation, enzyme-inhibition experiments, and comparisons with cancer-type-matched nonmalignant cells. Such validation is essential to establish the biological relevance, therapeutic selectivity, and clinical potential of the identified biomarkers and metabolic targets.

## Abbreviations

**symbol cancer type**

ACC: adrenocortical carcinoma
BLCA: bladder urothelial carcinoma
BRCA: breast invasive carcinoma
CESC: cervical squamous cell carcinoma and endocervical adenocarcinoma
CHOL: cholangiocarcinoma
COAD: colon adenocarcinoma
DLBC: diffuse large B-cell lymphoma
ESCA: esophageal carcinoma
GBM: glioblastoma multiforme
HNSC: head and neck squamous cell carcinoma
KICH: kidney chromophobe
KIRC: kidney renal clear cell carcinoma
KIRP: kidney renal papillary cell carcinoma
LAML: acute myeloid leukemia
LGG: brain lower grade glioma
LIHC: liver hepatocellular carcinoma
LUAD: lung adenocarcinoma
LUSC: lung squamous cell carcinoma
MESO: mesothelioma
OV: ovarian serous cystadenocarcinoma
PAAD: pancreatic adenocarcinoma
PCPG: pheochromocytoma and paraganglioma
PRAD: prostate adenocarcinoma
READ: rectum adenocarcinoma
SARC: sarcoma
SKCM: skin cutaneous melanoma
STAD: stomach adenocarcinoma
TGCT: testicular germ cell tumors
THCA: thyroid carcinoma
THYM: thymoma
UCEC: uterine corpus endometrial carcinoma
UCS: uterine carcinosarcoma
UVM: uveal melanoma

## Acknowledgements

This work was supported by the National Science and Technology Council, Taiwan (grant nos. NSTC113-2221-E-194-001 and NSTC113-2321-B-030-001).

## Data accessibility

The datasets supporting this article are available as part of the electronic supplementary material. The optimization code used in this study is available from GitHub: https://github.com/chmfsw/ACTD-toolbox/tree/main

## Authors’ contributions

F.S.W. conceived and designed the study and contributed to drafting the manuscript; S.Y.P. and Y.F.Y. performed model reconstruction and data analysis; K.T.H. coordinated the study and contributed to manuscript drafting. All authors gave final approval for publication and agree to be held accountable for the work.

## Competing interests

The authors declare no competing interests.

## Notes

### Competing Interest Statement

The authors have declared no competing interest.

