## Supplementary materials for "Constraint-Based Pan-Cancer Metabolic Modeling Identifies Medium-Robust Biomarkers and Broad-Spectrum Anticancer Targets"

Chiayi 621301, Taiwan

**Abstract**

**Background:** Cancer metabolic reprogramming varies across tumour types and nutrient environments, complicating the identification of robust metabolic biomarkers and therapeutic targets. We developed a constraint-based pan-cancer framework to identify medium-robust metabolic alterations and broadly effective candidate enzyme targets.

**Methods:** Transcriptomic profiles from 33 TCGA cancer types were integrated with Recon3D to reconstruct cancer- and healthy-tissue-specific genome-scale metabolic models. Parsimonious multi-objective metabolite flow variability analysis was performed under five nutrient conditions to classify differential rates of metabolite synthesis, uptake, and secretion. An extended Anticancer Target Discovery framework incorporating fuzzy multi-objective optimization was employed to evaluate candidate enzyme perturbations based on cancer-cell inhibition, ATP suppression, preservation of healthy-tissue metabolism, and robustness across nutrient conditions.

**Findings:** The models predicted recurrent increases in cardiolipin-related lipid synthesis and aromatic amino acid metabolism, enhanced uptake of essential amino acids and choline, and elevated lactate secretion across multiple cancer–medium combinations. CRLS1 emerged as the most broadly effective predicted enzyme target, satisfying the treatment criteria across all 165 cancer–medium combinations, whereas PGS1 and PTPMT1 exhibited broad but incomplete coverage. These findings highlight mitochondrial phospholipid metabolism as a potential shared vulnerability across cancer types and provide testable candidates for metabolomic and experimental validation.

**Keywords:** Computational systems biology; cancer metabolism; flux balance analysis; hybrid differential evolution; fuzzy multi-objective optimization; anticancer target discovery.

**Supplementary Materials**

**S1. Reconstruction of pan-cancer metabolic models**

CA and HT genome-scale metabolic models were reconstructed with CORDA for 33 TCGA cancer types. Table S1 gives the size of each model.

**Table S1. Numbers of metabolites and reactions in CORDA-reconstructed cancer-associated and healthy-tissue genome-scale metabolic models for 33 TCGA cancer types.**

*Legend:* Each entry is given as (Spc, Rxn), the numbers of metabolite species and reactions retained in the CA and HT model of each cancer type. Types marked † (ACC, DLBC, LAML, LGG, MESO, OV, SKCM, TGCT, UCS and UVM) use the generic Recon3Dmodel (5,835 metabolites, 10,600 reactions) as the HT reference, because matched adjacent-normal samples were unavailable for nine of them (ACC, DLBC, LAML, LGG, MESO, OV, TGCT, UCS and UVM) and SKCM had only one normal sample. Comparisons involving these ten types may therefore reflect differences in reference-model composition and network coverage, as well as cancer-associated metabolic alterations.

| **Type** | $\left( \boldsymbol{Spc, Rxn} \right)_{\boldsymbol{CA}}$ | $\left( \boldsymbol{Spc, Rxn} \right)_{\boldsymbol{HT}}$ | **Type** | $\left( \boldsymbol{Spc,Rxn} \right)_{\boldsymbol{CA}}$ | $\left( \boldsymbol{Spc,Rxn} \right)_{\boldsymbol{HT}}$ |
| --- | --- | --- | --- | --- | --- |
| ACC^†^ | (3669, 5963) | (5835, 10600) | LUSC | (3667, 5974) | (4283, 6815) |
| BLCA | (3732, 5896) | (3758, 6021) | MESO^†^ | (3726, 6115) | (5835, 10600) |
| BRCA | (3644, 5907) | (3830, 5917) | OV^†^ | (3662, 6026) | (5835, 10600) |
| CESC | (3617, 5823) | (3828, 5919) | PAAD | (3820, 6252) | (3851, 6001) |
| CHOL | (3905, 6155) | (4113, 6456) | PCPG | (3672, 5911) | (4470, 7062) |
| COAD | (3807, 6049) | (3667, 5890) | PRAD | (3684, 5925) | (4252, 6775) |
| DLBC^†^ | (3725, 6078) | (5835, 10600) | READ | (3881, 6296) | (3730, 5919) |
| ESCA | (3728, 6121) | (3866, 6178) | SARC | (3688, 6107) | (3787, 6015) |
| GBM | (3704, 6079) | (3798, 6072) | SKCM^†^ | (3625, 5820) | (5835, 10600) |
| HNSC | (3695, 6070) | (3819, 6133) | STAD | (3883, 6353) | (3643, 5874) |
| KICH | (3767, 6082) | (3640, 5599) | TGCT^†^ | (3698, 6078) | (5835, 10600) |
| KIRC | (3874, 6259) | (4229, 6444) | THCA | (3736, 5782) | (3703, 5743) |
| KIRP | (3847, 6234) | (3837, 5891) | THYM | (3731, 6094) | (4271, 6517) |
| LAML^†^ | (4022, 6359) | (5835, 10600) | UCEC | (3697, 5936) | (3700, 5608) |
| LGG^†^ | (3819, 6296) | (5835, 10600) | UCS^†^ | (3638, 5837) | (5835, 10600) |
| LIHC | (4014, 6361) | (4183, 6689) | UVM^†^ | (3819, 6257) | (5835, 10600) |
| LUAD | (3805, 6202) | (4262, 6753) |  |  |  |

**S2. Classification of differential metabolite-flow ranges under the reference configuration (Case 0)**

Table S2 reports, for each cancer type, the number of intracellular metabolites whose synthesis-rate classification was identical under all five nutrient conditions. Case 0 is the reference formulation (biomass and ATP production maximised in CA models; expression-dependent UFD reaction weights; see Table S3).

**Table S2. Consistently classified intracellular metabolite synthesis rates per cancer type under Case 0 (reference configuration: biomass and ATP objectives; expression-dependent UFD weights).**

*Legend:* Each entry is the number of intracellular metabolites assigned to the indicated category in the TCGA-derived CA models relative to the corresponding HT models. A metabolite is counted in a category only if that classification is maintained under all five nutrient conditions. Categories are CI (+3), PI (+2), II (+1), ID (−1), PD (−2) and CD (−3); the zero-score category (UC, 0) is not shown. Cancer types are sorted by decreasing CI count. Counts depend on model reconstruction, objective functions, nutrient constraints and the choice of HT reference (see Table S1).

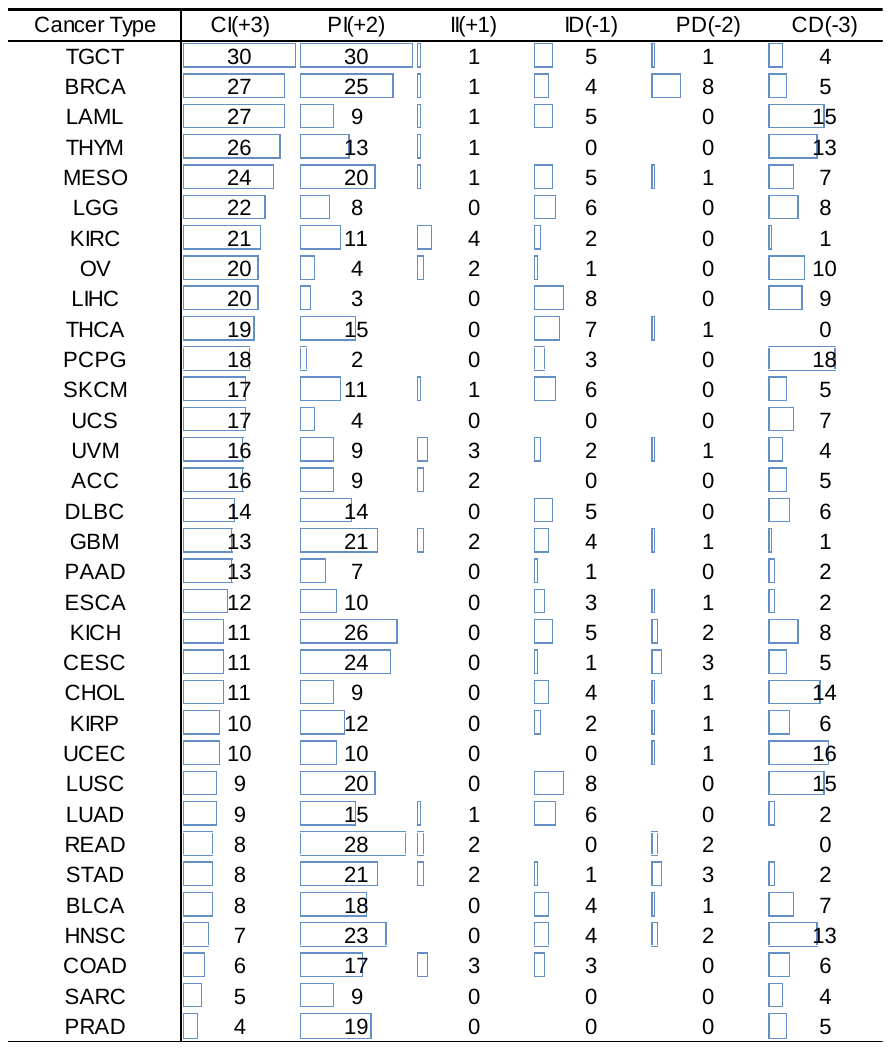

In most cancer models, positive classifications (CI, PI and II) outnumbered negative ones (ID, PD and CD); PCPG was the only exception (20 positive vs 21 negative). TGCT, BRCA, LAML and THYM had the largest numbers of CI features (26–30), whereas PCPG (18), UCEC (16), LAML (15) and LUSC (15) had the largest numbers of CD features. Thus, increased feasible metabolite-production capacity predominated in many reconstructed cancer models, although the balance between increased and decreased synthesis varied considerably among cancer types.

This predominance should not be read as universal metabolic activation. The counts depend on reconstruction, objective-function choice, network coverage, nutrient constraints and reference-model composition. In particular, comparisons against genericRecon3D rather than a matched tissue-specific HT model may partly reflect differences in network structure. The counts therefore describe the behaviour of the reconstructed models under the stated assumptions, not measured global metabolic activity in tumours.

CI and CD give the strongest model-based discrimination because the predicted CA and HT flux ranges do not overlap, whereas PI and PD involve overlapping ranges and discriminate less well. In every case the classifications describe differences in feasible metabolic ranges, not experimentally measured distributions in patient samples. The identified features should be regarded as model-predicted candidate metabolic biomarkers that require experimental and clinical validation.

**S3. Sensitivity analysis of pMOMFVA predictions**

**S3.1. Computational configurations**

To test how sensitive pMOMFVA predictions are to modelling assumptions, four configurations were evaluated across the 33 TCGA-derived cancer types and five nutrient conditions. They differ in the objective functions assigned to CA models and in reaction weights used in the UFD stage; the UFD weighting scheme was applied to both CA and HT models (Table S3).

**Table S3. Computational configurations (Cases 0–3) used to test the sensitivity of pMOMFVA predictions to objective-function selection and UFD reaction weighting**

*Legend:* All configurations were run for 33 cancer types and five nutrient conditions. Case0 is the reference formulation used in the main manuscript. Case 1 changes only the reaction weighting, Case 2 changes only the objective function, and Case 3 changes both. The HT-model objective is ATP production in all cases.

| Case | CA-model objective functions | UFD reaction weights (CA and HT) | Purpose |
| --- | --- | --- | --- |
| 0 | Maximize biomass and ATP production | Expression-dependent | Reference configuration (main manuscript) |
| 1 | Maximize biomass and ATP production | Uniform | Effect of reaction weighting |
| 2 | Maximize biomass production only | Expression-dependent | Effect of objective selection |
| 3 | Maximize biomass production only | Uniform | Combined alternative settings |

The expression-dependent reaction weights are defined in Equation (2) of the main manuscript, whereas the uniform weighting scheme assigns a weight of 1 to all reactions.

**S3.2. Influence of objective-function selection and reaction weighting**

Both the CA-model objective functions and the UFD reaction weights affected the classification patterns (Table S4). Summed over the 33 cancer types, biomass-only optimisation (Cases 2 and 3) shifted assignments strongly towards CI (1,663 and 1,548) and almost eliminated PI (5 and 4). The combined biomass and ATP objectives (Cases 0 and 1) gave more differentiated profiles (CI 489 and 747; PI 476 and 199). Objective selection therefore strongly influences the predicted separation between CA and HT metabolite-flow ranges.

The effect of reaction weighting depended on the objective. Under the combined objectives, replacing expression-dependent with uniform weights (Case 0 vs Case 1) raised the CI count from 489 to 747 and lowered the PI count from 476 to 199 and the CD count from 225 to 158. Under the biomass-only objective (Case 2 vs Case 3), CI changed less (1,663 to 1,548), whereas ID rose from 100 to 163 and CD fell from 232 to 188. Per-cancer counts are given in Table S2 (Case 0) and Tables S5–S7 (Cases 1–3).

**Table S4. Aggregate counts of consistently classified intracellular metabolite synthesis rates across 33 cancer types under Cases 0–3.**

*Legend:* Values are the sums, over the 33 cancer types, of the within-cancer counts of metabolites assigned to the same category under all five nutrient conditions. They are not numbers of unique metabolites pooled across cancers. Categories are CI (+3), PI (+2), II(+1), ID (−1), PD (−2) and CD (−3). Cases 0–3 are defined in Table S3.

| Case | CI(+3) | PI(+2) | II(+1) | ID(-1) | PD(-2) | CD(-3) |
| --- | --- | --- | --- | --- | --- | --- |
| 0 | 489 | 476 | 27 | 105 | 30 | 225 |
| 1 | 747 | 199 | 19 | 135 | 38 | 158 |
| 2 | 1663 | 5 | 4 | 100 | 31 | 232 |
| 3 | 1548 | 4 | 4 | 163 | 28 | 188 |

**Table S5. Consistently classified intracellular metabolite synthesis rates per cancer type under Case 1 (biomass and ATP objectives; uniform UFD weights).**

*Legend:* Each entry is the number of intracellular metabolites assigned to the indicated category under all five nutrient conditions. Categories are CI (+3), PI (+2), II (+1), ID (−1),PD (−2) and CD (−3); UC (0) is not shown. Cancer types are sorted by decreasing CI count.

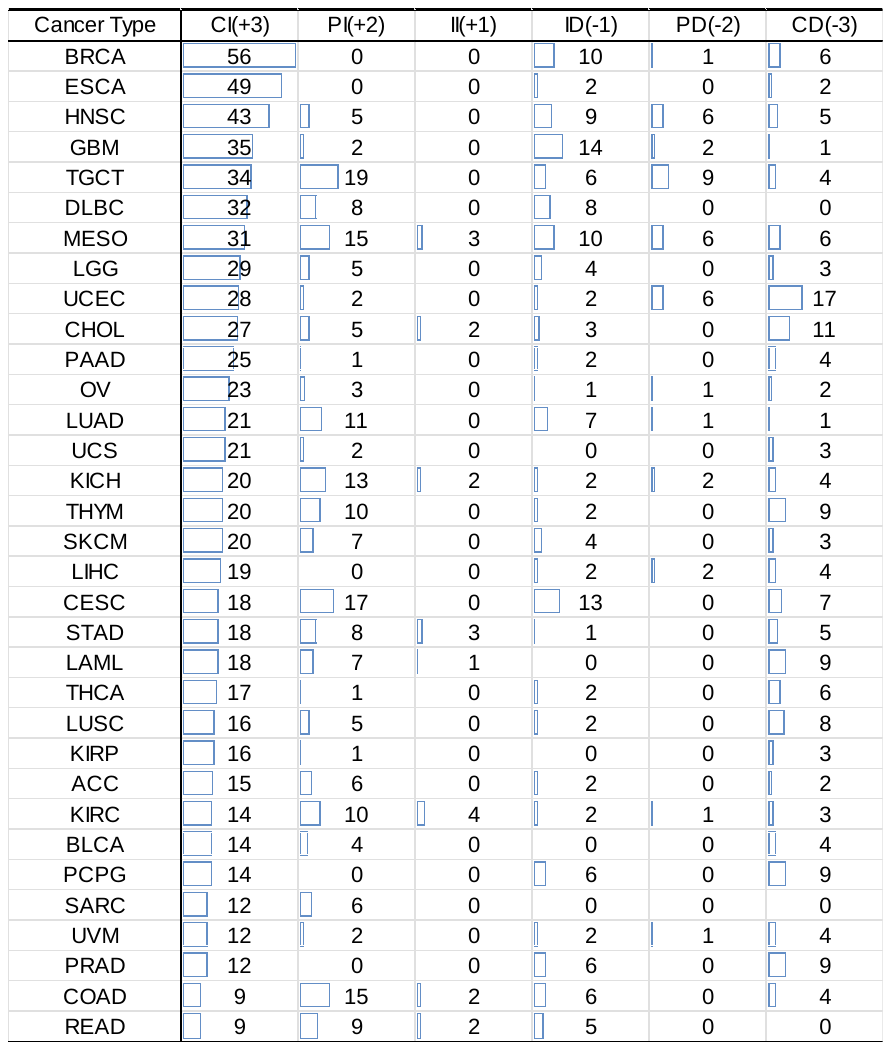

**Table S6. Consistently classified intracellular metabolite synthesis rates per cancer type under Case 2 (biomass-only objective; expression-dependent UFD weights).**

*Legend:* As for Table S5, for Case 2.

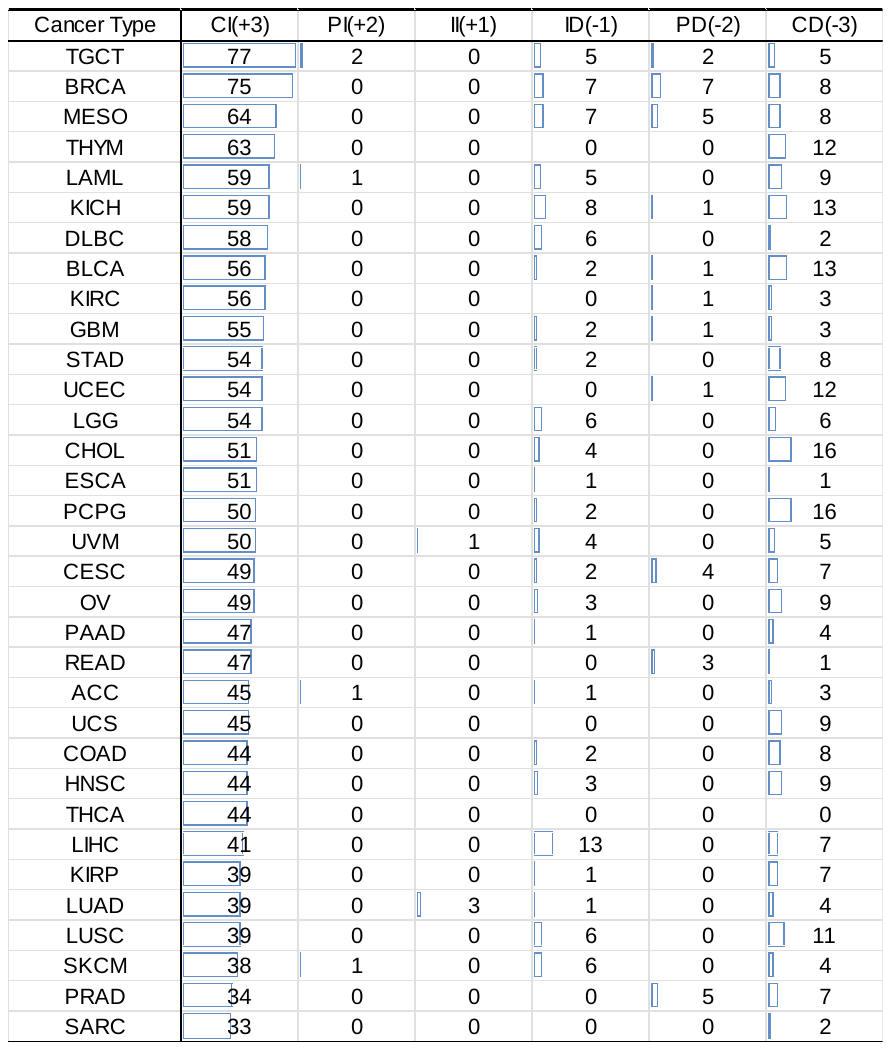

**Table S7. Consistently classified intracellular metabolite synthesis rates per cancer type under Case 3 (biomass-only objective; uniform UFD weights).**

*Legend:* As for Table S5, for Case 3.

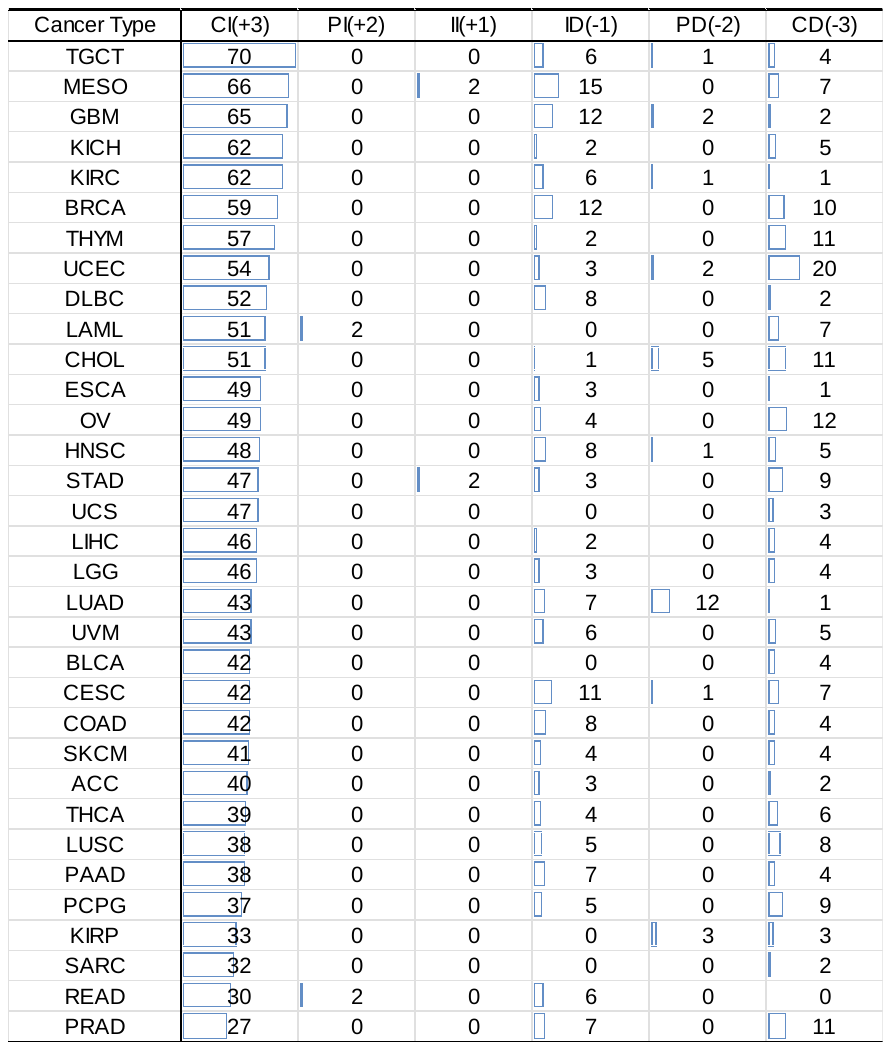

**S3.3. Sensitivity of intracellular synthesis and exchange predictions**

We next compared classification counts and average scores for selected intracellular metabolite synthesis rates, secretion and uptake across all 165 cancer–medium instances (33 cancer types × five nutrient conditions). Reference (Case 0) results are in Table 1 of the main manuscript; Cases 1–3 are in Tables S8–S10.

**Table S8. Classification counts and average scores for selected intracellular synthesis, secretion and uptake rates under Case 1 (biomass and ATP objectives; uniform UFD weights).**

*Legend:* Each row summarises 165 cancer–medium instances. The predicted CA–HT difference in each instance is classified as CI (+3), PI (+2), II (+1), UC (0), ID (−1), PD (−2) or D (−3). The average score (Ave.) is the arithmetic mean of these scores over the 165instances; it is neither a fold change nor a measured flux. Rows are grouped by rate type intracellular synthesis, secretion, uptake). Pravastatin-derived entries are xenobiotic-associated features.

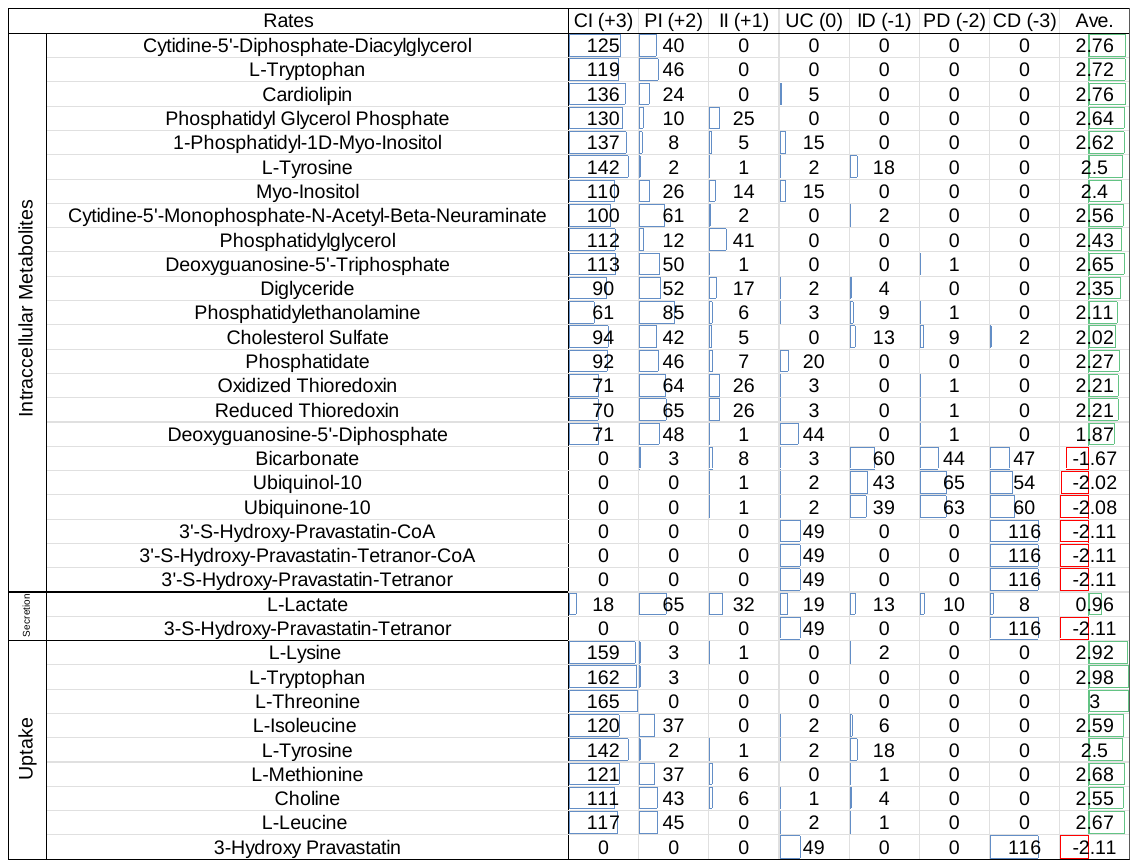

**Table S9. Classification counts and average scores for selected intracellular synthesis, secretion and uptake rates under Case 2 (biomass-only objective; expression-dependent UFD weights).**

*Legend:* As for Table S8, for Case 2.

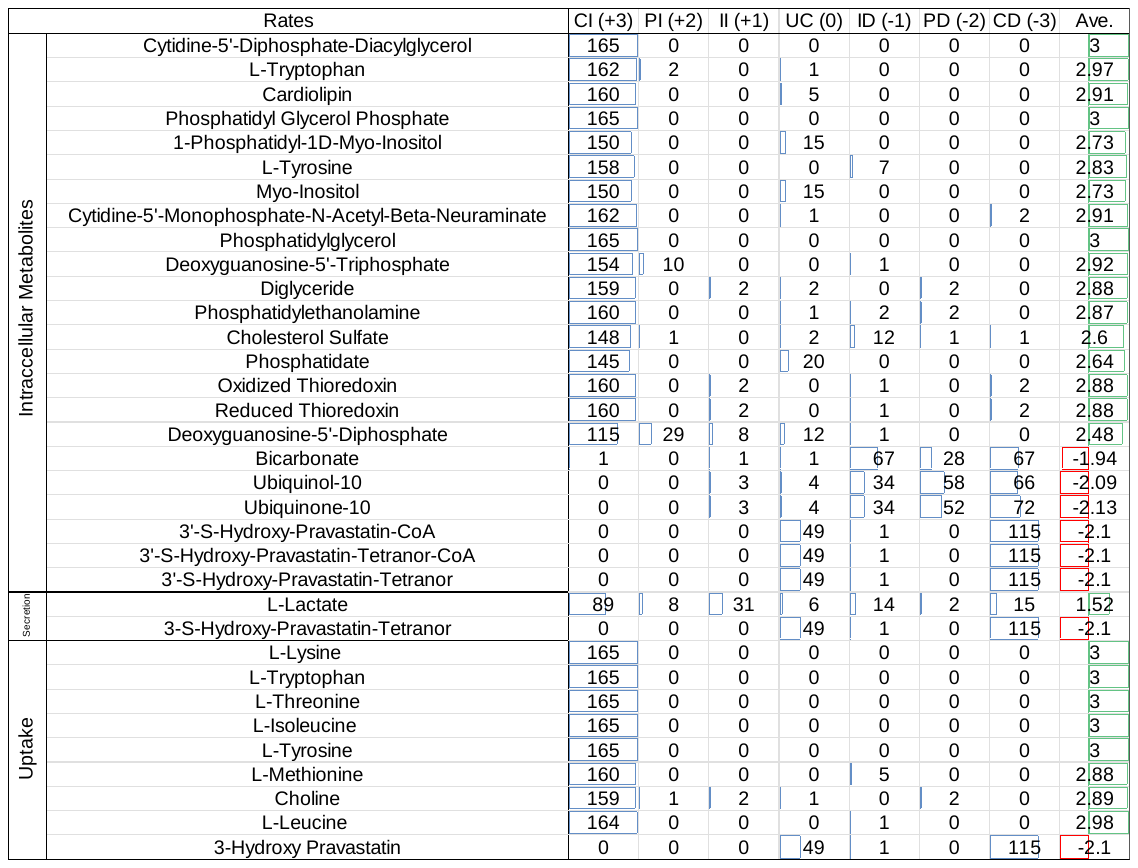

**Table S10. Classification counts and average scores for selected intracellular synthesis, secretion and uptake rates under Case 3 (biomass-only objective; uniform UFD weights).**

*Legend:* As for Table S8, for Case 3.

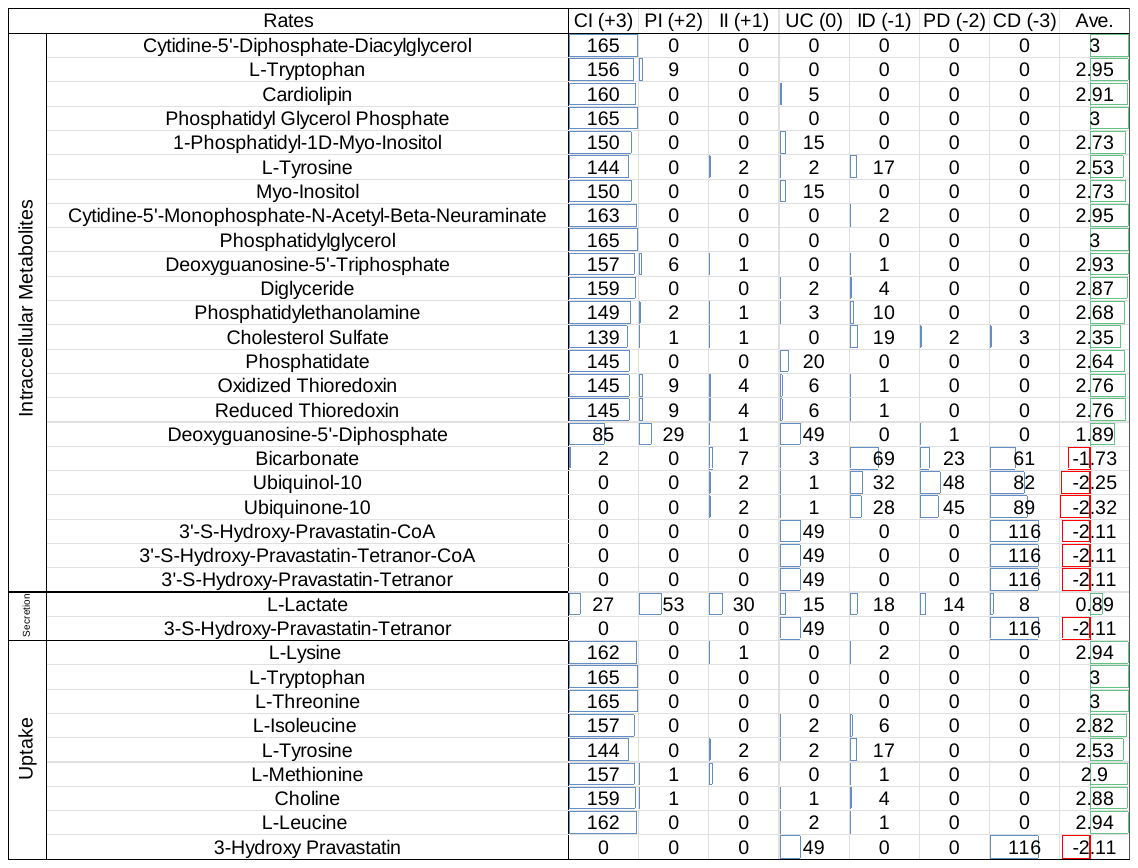

Several predictions kept the same direction across all four configurations. Intracellular synthesis of CDP-diacylglycerol, cardiolipin, L-tryptophan and phosphatidylglycerol phosphate always had positive average scores (2.76–3.00, 2.75–2.91, 2.72–2.97 and 2.59–3.00, respectively), whereas bicarbonate (−1.94 to −1.67), ubiquinol-10 (−2.25 to −1.92) andubiquinone-10 (−2.32 to −1.93) always had negative average scores. Pravastatin-associated intracellular and exchange features also kept negative average scores (about−2.08 to −2.11), but these are xenobiotic-associated predictions and 49 cancer–medium instances were scored zero (UC) in every configuration.

Amino acid and choline uptake was predominantly positive in all configurations, particularly for L-lysine, L-tryptophan, L-threonine, L-isoleucine and choline, although the size and spread of the scores differed among metabolites. L-Lactate secretion was the most sensitive to the modelling assumptions: its average score ranged from 0.89 to 1.52and its classifications spanned all seven categories. Overall, a subset of predictions is directionally robust, whereas others depend more on the objective functions and reaction weights.

**S3.4. Overall interpretation**

The sensitivity analysis supports two conclusions. First, several prominent alterations remained directionally consistent in all four configurations: increased cardiolipin-related synthesis, enhanced amino acid uptake, and reduced ubiquinone- and ubiquinol-associated synthesis. Second, biomass-only optimisation produced a much larger proportion of CI classifications and a more concentrated distribution of outcomes than the combined biomass and ATP objectives.

The reference configuration (Case 0) gives a more differentiated basis for comparing alterations across cancer types and nutrient conditions. Greater differentiation does not, however, imply greater biological accuracy, and experimental validation is needed to determine which objective formulation best represents the metabolism of individual cancers.

**S4. Sensitivity analysis of predicted CRLS1 inhibition**

To test the robustness of CRLS1 as a candidate anticancer target, its predicted inhibitory effects were compared across Cases 0–3. Case 0 results are shown in Figure 4 of the main manuscript and Cases 1–3 in Figures S1–S3. Two scores are shown for each cancer type. The cancer-cell inhibition score reflects the predicted suppression of the CA-model objectives (growth and ATP production in Cases 0 and 1; biomass production in Cases 2and 3); higher values mean stronger suppression. The metabolic-deviation score reflects the predicted metabolic response relative to the CA and HT reference templates; higher values mean a more favourable response. In all figures, the blue segment is the minimum score across the five nutrient conditions and the orange segment is the maximum minus the minimum, so the total bar height is the maximum score and the orange segment shows nutrient-dependent variability.

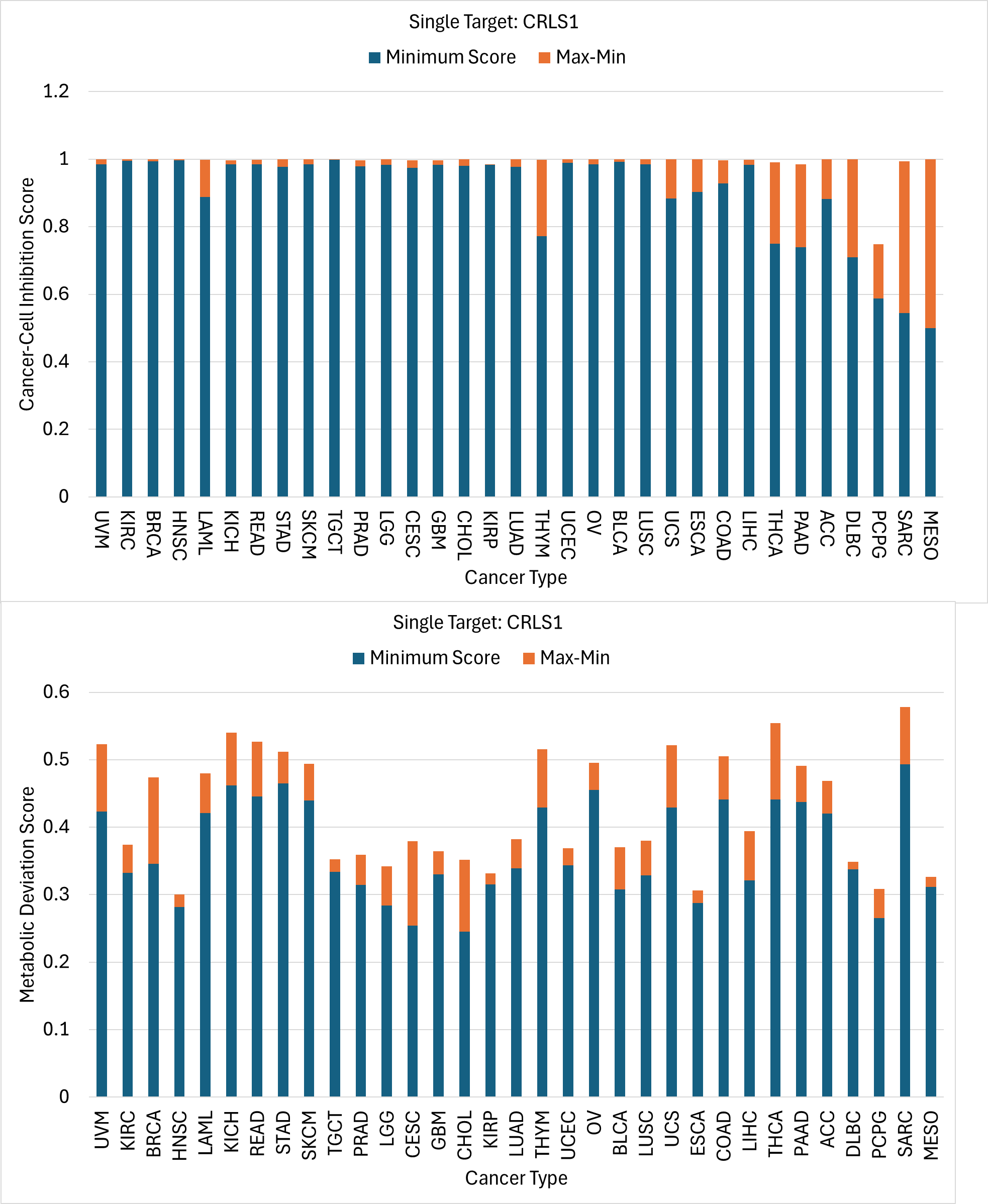

**Figure S1. Predicted cancer-cell inhibition (upper) and metabolic-deviation (lower) scores after CRLS1 inhibition under Case 1 (biomass and ATP objectives; uniform UFD weights).**

*Legend:* Scores are shown for 33 cancer types, each under five nutrient conditions. Blue segments show the minimum score across the five media; orange segments show the maximum minus the minimum. Cancer types are ordered as in Figure 4.

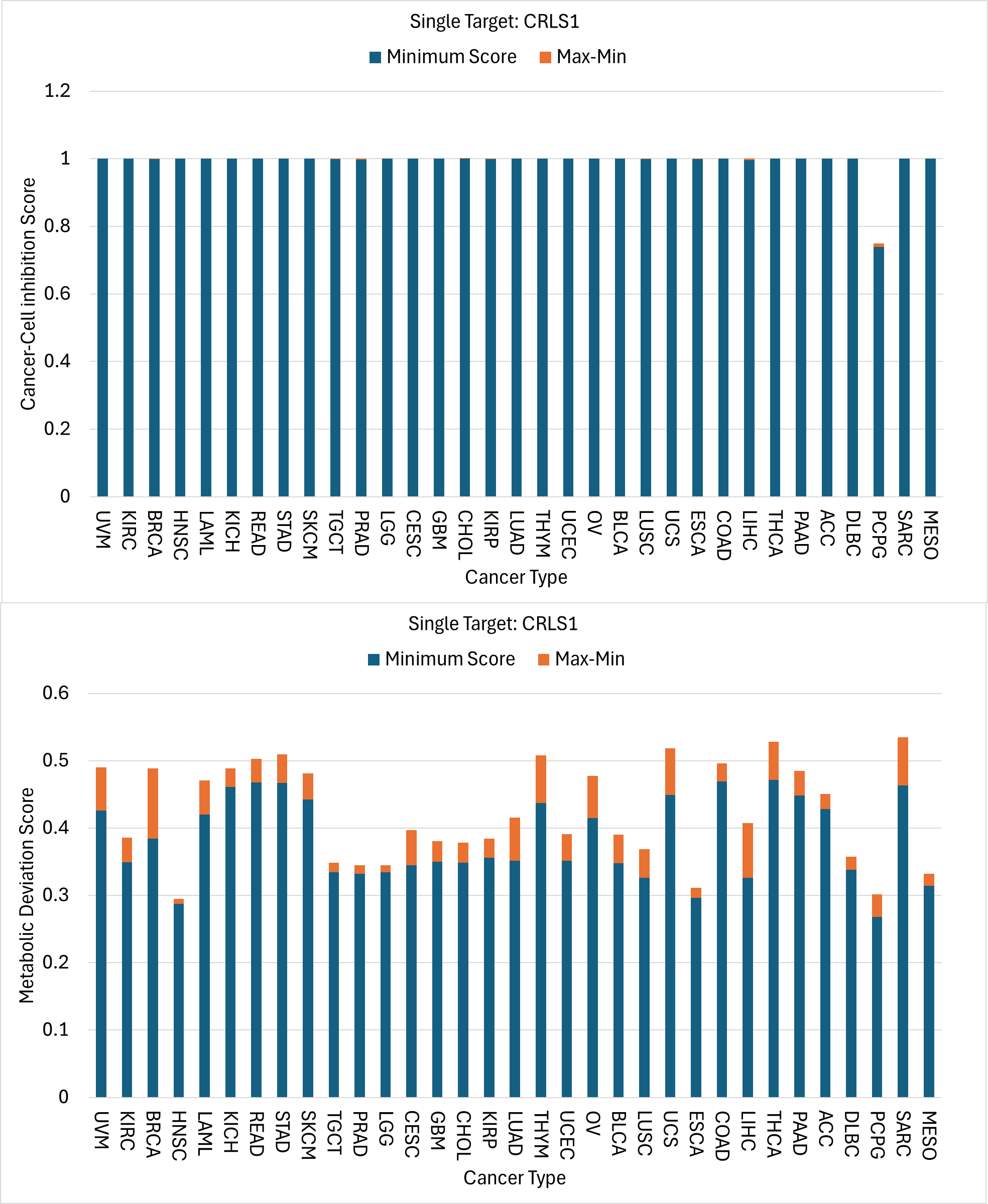

**Figure S2. Predicted cancer-cell inhibition (upper) and metabolic-deviation (lower)scores after CRLS1 inhibition under Case 2 (biomass-only objective; expression-dependent UFD weights).**

*Legend:* As for Figure S1, for Case 2. Inhibition scores approach 1 for all cancer types except PCPG (minimum ≈ 0.75).

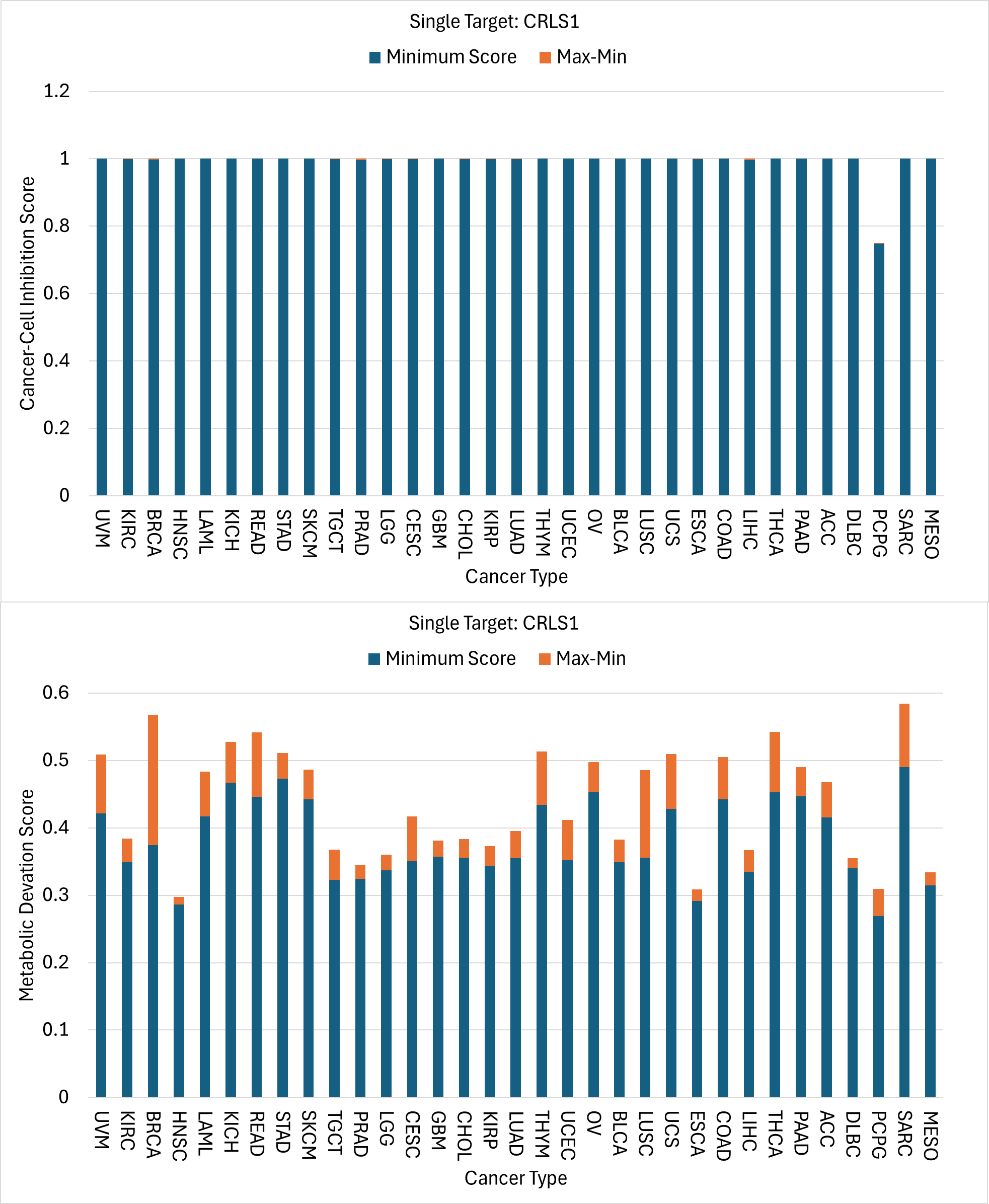

**Figure S3. Predicted cancer-cell inhibition (upper) and metabolic-deviation (lower)scores after CRLS1 inhibition under Case 3 (biomass-only objective; uniform UFD weights).**

*Legend:* As for Figure S1, for Case 3. Inhibition scores approach 1 for all cancer types except PCPG (minimum ≈ 0.75).

**S4.1 Comparison across configurations**

CRLS1 inhibition gave generally high cancer-cell inhibition scores in all four configurations, although their magnitude and nutrient-dependent variability differed. Under Case 0, minimum inhibition scores were about 0.5–0.6 for several cancer types, whereas many others approached unity. Case 1 gave minimum scores close to unity for most cancer types, with a few lower exceptions (for example, MESO, SARC and PCPG). Under biomass-only optimisation (Cases 2 and 3), minimum scores approached unity for nearly all cancer types, with PCPG remaining the notable exception (about 0.75).

In contrast, metabolic-deviation scores remained heterogeneous across cancer types and media, mostly within about 0.25–0.60, with broadly similar patterns in all four configurations. CRLS1 inhibition therefore retained broad predicted anticancer activity, but the magnitude and nutrient-dependent variability of its effects were more sensitive to the choice of cancer-cell objective than to the UFD weighting scheme. The heterogeneity of the deviation scores also shows that strong predicted inhibition of cancer cells does not necessarily accompany an equally favourable overall metabolic response.

**S5. Genetic-dependency analysis of predicted anticancer targets**

To assess the biological relevance of the predicted targets, DepMap genetic-dependency data were examined for three cardiolipin-biosynthesis enzymes across ten cancer types: PGS1 (synthesis of phosphatidylglycerol phosphate), PTPMT1 (its dephosphorylation to phosphatidylglycerol) and CRLS1 (the final step to cardiolipin).

**Table S11. Frequency of genetic dependency on CRLS1, PGS1 and PTPMT1 across 582 DepMap cancer cell lines from ten cancer types.**

*Legend:* Each entry is the number of cell lines classified as dependent on the indicated gene divided by the number of cell lines evaluated for that cancer type (n/N). Dependency was defined from genome-wide CRISPR knockout screens in DepMap 24Q2 as a gene-effect score ≤ −0.5. The total row gives the overall counts and percentages across all ten cancer types. These data provide experimental context for the predicted metabolic vulnerabilities but do not measure therapeutic selectivity or the efficacy of pharmacological enzyme inhibition.

| Cancer Type | CRLS1 | PGS1 | PTPMT1 |
| --- | --- | --- | --- |
| BRCA | 24/53 | 27/53 | 23/53 |
| COAD | 24/48 | 25/48 | 18/48 |
| DLBC | 47/93 | 68/93 | 51/93 |
| HNSC | 44/77 | 55/77 | 45/77 |
| LIHC | 11/24 | 18/24 | 14/24 |
| LUAD | 71/126 | 68/126 | 60/126 |
| OV | 29/59 | 30/59 | 32/59 |
| PAAD | 20/47 | 25/47 | 24/47 |
| PRAD | 3/10 | 5/10 | 4/10 |
| STAD | 23/45 | 29/45 | 23/45 |
| Total | 296/582 (50.9%) | 350/582 (60.1%) | 294/582 (50.5%) |

Across the 582 cell lines, dependencies on CRLS1, PGS1 and PTPMT1 were found in 296 (50.9%), 350 (60.1%) and 294 (50.5%) lines, respectively. PGS1 dependency was more frequent than CRLS1 dependency in nine of the ten cancer types (LUAD was the exception), even though CRLS1 had the broader predicted effect in the constraint-based simulations. This discrepancy shows that experimentally observed genetic dependencies and model-predicted metabolic vulnerabilities can differ, possibly because of cellular context, network constraints, nutrient availability, and differences between genetic perturbation and simulated enzyme inhibition.

Nevertheless, the DepMap data provide supportive experimental evidence that mitochondrial cardiolipin biosynthesis is a potential metabolic vulnerability in cancer. Further experiments are needed to determine whether perturbing these enzymes can selectively inhibit cancer cells while preserving essential metabolic functions in non-malignant tissues.
